# Single-Cell Resolution of Individual Variation in Hypothalamic Neurons Allows Targeted Manipulation Affecting Social Motivation

**DOI:** 10.1101/2025.03.10.642464

**Authors:** S. Sarafinovska, S.K. Koester, L.Z. Fang, J.W. Thorpe, S.M. Chaturvedi, J. Ji, E.F. Jones, D. Selmanovic, D.J. Kornbluth, M.R. Barrett, G.M. Rurak, S.E. Maloney, M.C. Creed, R.D. Mitra, J.D. Dougherty

**Affiliations:** Department of Genetics, Washington University School of Medicine, Saint Louis, MO, USA; Department of Psychiatry, Washington University School of Medicine, Saint Louis, MO, USA; Washington University Pain Center, Department of Anesthesiology, St. Louis, MO, USA; Intellectual and Developmental Disabilities Research Center, Washington University School of Medicine, St. Louis, MO, 63110-1093, USA; Edison Family Center for Genome Sciences and Systems Biology, Washington University School of Medicine, Saint Louis, MO, USA

## Abstract

Despite decades of research, connecting molecular and cellular phenotypes to complex behavioral traits remains an elusive goal^1^. Social motivation exhibits individual trait variation^2^, which we hypothesize is mediated by molecular and cellular variability across hypothalamic neurons. To test this, we generated single-nucleus RNA-sequencing profiles^3,4^ of >120,000 neurons from tuberal hypothalamus and adjacent thalamus in 36 mice, balanced across sex and autism-associated mutation^5^, with all mice assessed for social motivation^2^. First, we show that molecular activation patterns predict behavior across individuals: specifically, activation of paraventricular *Agtr1a*+ (angiotensin receptor 1a) neurons predicted reduced social behavior. Subsequent inhibition of AGTR1A with telmisartan—an FDA-approved antihypertensive^6^—improved social orienting. Second, we show natural variation in neuronal proportions—likely arising from stochastic developmental events^7^—is sufficient to shape adult behavior even among genetically-identical individuals: we identified multiple neuronal populations whose relative abundance predicted social reward-seeking behavior. Chemogenetic inhibition of one such population, *Nxph4*+ neurons of the postero-lateral hypothalamus^8^, suppressed multiple aspects of social motivation. This work establishes proof-of-principle for an approach where single-cell genomics precisely maps neural substrates governing behavior. This approach revealed that stochastic variations in neuronal architecture deterministically influence social motivation, and enabled identification of therapeutically-actionable targets with immediate translational potential for disorders with social deficits.

## INTRODUCTION

One great challenge in neuroscience is how to connect molecular and cellular phenotypes to their behavioral consequences^1^, as gene mutations, sex, and uncharacterized individual differences all determine behavior^9^. Since genes function at the molecular level to influence cellular function yet also impact behavior, novel methods are needed to bridge this gap and define the cellular substrates that drive behavioral differences. As markers of neuronal activity, immediate early genes (IEGs), like *Fos,* have been used to anatomically define regions activated by a range of behaviors^10–13^. More recently, Act-seq and related methods enabled faithful mapping of such activity-induced gene expression to cell types within a region using single-nucleus RNA sequencing (snRNAseq)^14,15^. For example, single-cell neuronal activity, as measured by IEG expression, identified cell types activated in a fasting state^16^ or neurons activated in social behaviors including aggression, mating, and parenting^17,18^. These studies performed binary comparisons between groups: experimental vs. control animals. Single-neuronal IEG expression levels, however, could also serve as quantitative predictors of behavioral variation across individuals, achieving more sensitive associations between cell-type activity and behavioral outcomes. However, this would require large-scale multi-animal analyses^19^. Furthermore, IEG expression is but one source of individual cellular variation neglecting unmeasured differences in neuronal composition or baseline transcriptional states that could influence behavior.

Social motivation in rodents, measured through reward-seeking and orienting behaviors in social operant assays, is a stable trait that shows individual variation and is also influenced by sex and genetics^2,5,20^. Behavioral variation stems from internal drives potentially mediated by natural variation in neuronal composition and gene expression patterns^7,9,21–23^. Understanding cellular mediators of social variation has direct relevance for psychiatric conditions with core social deficits, including autism, depression, and schizophrenia^24–26^. Recent advances in single-cell profiling have enabled quantification of such variation across individuals, meaning we can now ask whether individual differences in cell type proportions and gene expression patterns predict variation in social behavior^19^. Understanding these cellular mediators of social variation could inform the development of targeted therapeutics for social deficits in psychiatric conditions^24–26^.

Here, we used snRNAseq to profile the tuberal hypothalamus and neighboring thalamus (hereafter, tuHY/TH) from 36 mice assessed for social motivation, balanced across sex and autism-associated genetic vulnerability (*Myt1l* genotype)^5^. Focusing on these well-mapped^16^, socially-relevant^17,18^, and sex-dimorphic^27,28^ regions, we developed an analytical framework treating both neuronal subtype proportions and IEG expression as continuous predictors of social behavior. This revealed complementary cellular mechanisms regulating social motivation: immediate-early gene expression analysis identified *Agtr1a*+ (angiotensin receptor 1a) neurons as therapeutically-actionable negative regulators of social motivation, with pharmacological blockade of angiotensin receptors enhancing social reward-seeking in wild-type mice. Novel analysis of cell type proportions revealed *Nxph4*+ neurons as positive regulators of social motivation which we confirmed through targeted chemogenetic inhibition. Overall, by leveraging natural variation across individuals, we established a new paradigm to associate cell types with behavior. Our findings reveal that natural variation in neuronal proportions—even among genetically-identical mice—is substantial enough to deterministically shape adult social behavior, suggesting that stochastic developmental events profoundly influence behavioral outcomes^7,21,23^. This approach identified novel neuronal regulators of social motivation and uncovered therapeutically-actionable targets, including a repurposable and well-tolerated FDA-approved drug^6,29^, with potential for treating social deficits in psychiatric disorders^24–26^.

## RESULTS

### Comprehensive tuberal hypothalamic and thalamic single-nucleus atlas from socially characterized mice

To identify the molecular factors that underlie an individual’s propensity for social motivation, we used single-cell combinatorial indexing^3,4^ to profile transcription from 138,033 nuclei from tuHY/TH in mice deeply characterized by a social operant task^2^. Thirty-six mice, split by sex and *Myt1l* genotype^5^, were conditioned to nose-poke for brief access to a sex- and age-matched novel stimulus animal (**Fig.1A**). Brains were harvested within 30 minutes after mice either mastered the task (“achievers,” showing 3 days of ≥40 nose-pokes, ≥75% accuracy, and ≥75% interaction attempts) or failed to meet these criteria (“non-achievers”) after 10 days of training^2^. Overall, female mice showed higher social motivation than males, as published^20^: females earned more rewards during training, spent more time investigating the other mouse, and had a higher proportion of achievers in the task (**Fig.1B,D,F, ED-Fig.1**), while *Myt1l* mutation did not reduce social motivation (**ED-Fig.1**). Yet, both sexes showed a wide range of variation in social behavior across genotypes (**Fig.1C,E,G**), indicating factors beyond sex or genotype also influence this trait.

**Figure 1:**
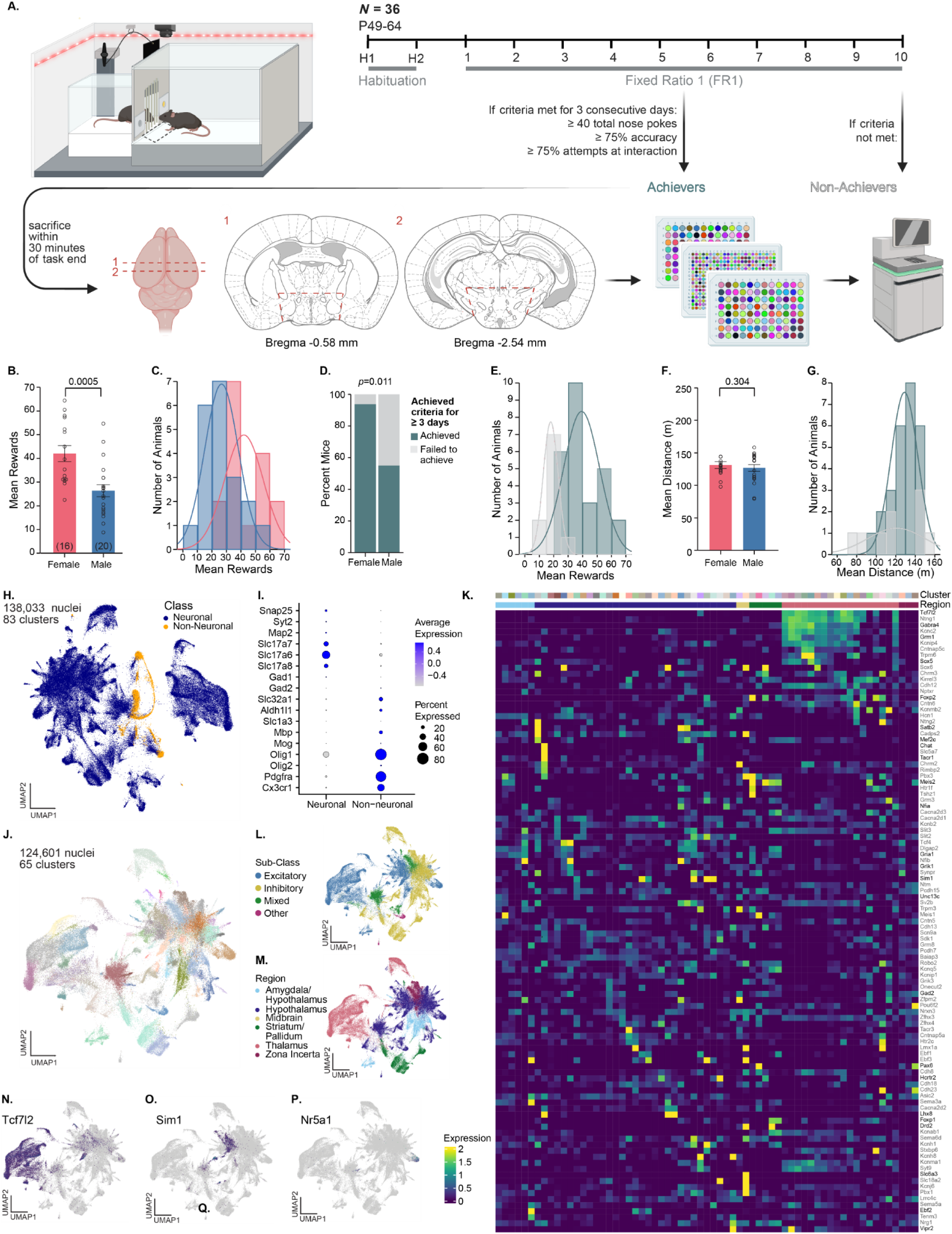
Comprehensive tuberal hypothalamic and neighboring thalamic (tuHY/TH) single-nucleus atlas from socially characterized mice. A. Schematic of the experimental setup showing the operant conditioning chamber where mice nose-poke to receive social rewards, followed by the timeline of the Fixed Ratio 1 (FR1) behavioral paradigm. Mice were classified as “Achievers” if they met performance criteria for three consecutive days (≥40 total nose-pokes, ≥75% accuracy, ≥75% attempts at interaction) or “Non-Achievers” if criteria were not met. Within 30 minutes of task completion, brains were harvested for analysis. Top-down brain diagram shows the locations of two coronal cuts (labeled 1 and 2), with corresponding coronal sections displayed, at Bregma -0.58 mm and Bregma -2.54 mm, respectively. Subsequent cut to collect tuHY/TH indicated in red dashed lines. Tissue was processed for single-nucleus combinatorial indexing RNA sequencing. B. Females earned significantly more social rewards on average across FR1 than males (p=0.0005; ANOVA), with a broad distribution of mean rewards for either sex (females in pink, males in blue; C.). D. Significant increase in proportion of mice that achieved criteria for ≥3 days versus those that failed to achieve criteria in females relative to males (p=0.011; chi-squared), with a broad spread of mean rewards by achiever status (achievers in teal, non-achievers in gray; E.). F. No differences in mean distance traveled by female versus male mice (p=0.304). G. Broad distribution of mean distance traveled across by achiever status (achievers in teal, non-achievers in gray). H. Uniform manifold approximation and projection (UMAP) visualization of full tuHY/TH dataset showing 138,033 nuclei clustered into 83 distinct cell populations, colored by neuronal (blue) versus non-neuronal (yellow) classification. I. Dot plot showing expression of marker genes differentiating neuronal from non-neuronal cell types. Dot size indicates the percent of cells expressing each gene, while color intensity represents the average expression level across all neuronal or non-neuronal clusters. J. UMAP of neuronal-only subset including 124,601 nuclei from tuHY/TH organized into 65 clusters. L. UMAP of neuronal subtypes classified as excitatory (blue), inhibitory (yellow), mixed (green), or other (magenta). M. UMAP of neurons organized by brain region, including amygdala/hypothalamus, hypothalamus, midbrain, striatum, pallidum, thalamus, and zona incerta. N. Expression patterns of thalamic marker, *Tcf7l2*. O. Expression patterns of paraventricular hypothalamic marker, *Sim1*. P. Expression patterns of the ventromedial hypothalamic marker, *Nr5a1*. K. Heatmap showing scaled expression of region-specific marker genes (rows) across hypothalamic and thalamic neuronal clusters (columns). Clusters are colored individually (bar, top) and by anatomical region (bar, bottom). Expression strength is indicated by color intensity as shown in the scale on the left (0-2). Key markers colored in black, including *Chat* (cholinergic), *Sim1* (paraventricular hypothalamus), *Hcrtr2* (orexin), *Slc6a3* (dopaminergic), and *Gad2* (GABAergic).

Across the tuHY/TH we identified 83 clusters, with neurons making up 91.8% and glia 8.2% (**Fig.1H-I**). Due to glial scarcity, we continued analysis in neuronal cells only, identifying 65 distinct neuronal subclusters (**Fig.1J-K**). Hypothalamic neurons were split between excitatory (glutamatergic) and inhibitory (GABAergic), while thalamic neurons were mostly excitatory (**Fig.1L**). Neuronal subtypes were largely transcriptionally distinguished by inferred regions, as observed^16^(**Fig.M**).

Next, we confirmed predicted annotations by manually examining marker gene expression. *Tcf7l2* was expressed broadly across the thalamic nuclei (**Fig.1N**). Universal markers of hypothalamic neurons do not exist, but we observed strong, distinct expressions of hypothalamic sub-region markers, including *Sim1* for the paraventricular nucleus and *Nr5a1* for the ventromedial hypothalamus (**Fig.1O-P**). Several populations showed mixed medial amygdala (MEA) and preoptic (PO) features by MapMyCells^30^ and preoptic features by HypoMap^16^, potentially due to shared origins, functionality, and transcriptional signatures^31^. Hence, we classified these as ‘PO-MEA’ neurons.

### Hypothalamus shows sex and genotype differences in cell proportion and gene expression

Next, we examined whether sex and genotype drive differences in the proportions of tuHY/TH neurons and their gene expression patterns. Nuclei were well-distributed across genotype (mean local inverse Simpson’s index (LISI)^32^ score = 1.97) and sex (mean LISI score = 1.91), with balanced sequencing depth of nuclei, genes, and unique molecular identifiers = (**Fig.2A-B,H-I**). Analysis of the established sex-differential marker *Moxd1*^28^ showed that males had significantly more *Moxd1*+ neurons, predominantly in cells predicted to be in the preoptic area (Lhx6/Lhx8 Gaba, PO-MEA-BST Mixed Gaba), potentially consistent with the sexually dimorphic nucleus (**Fig.2C**). Meanwhile, *Moxd1* expression levels per nucleus were similar between sexes, indicating *Moxd1* primarily marks sex-biased neuronal populations (**Fig.2C**).

**Figure 2:**
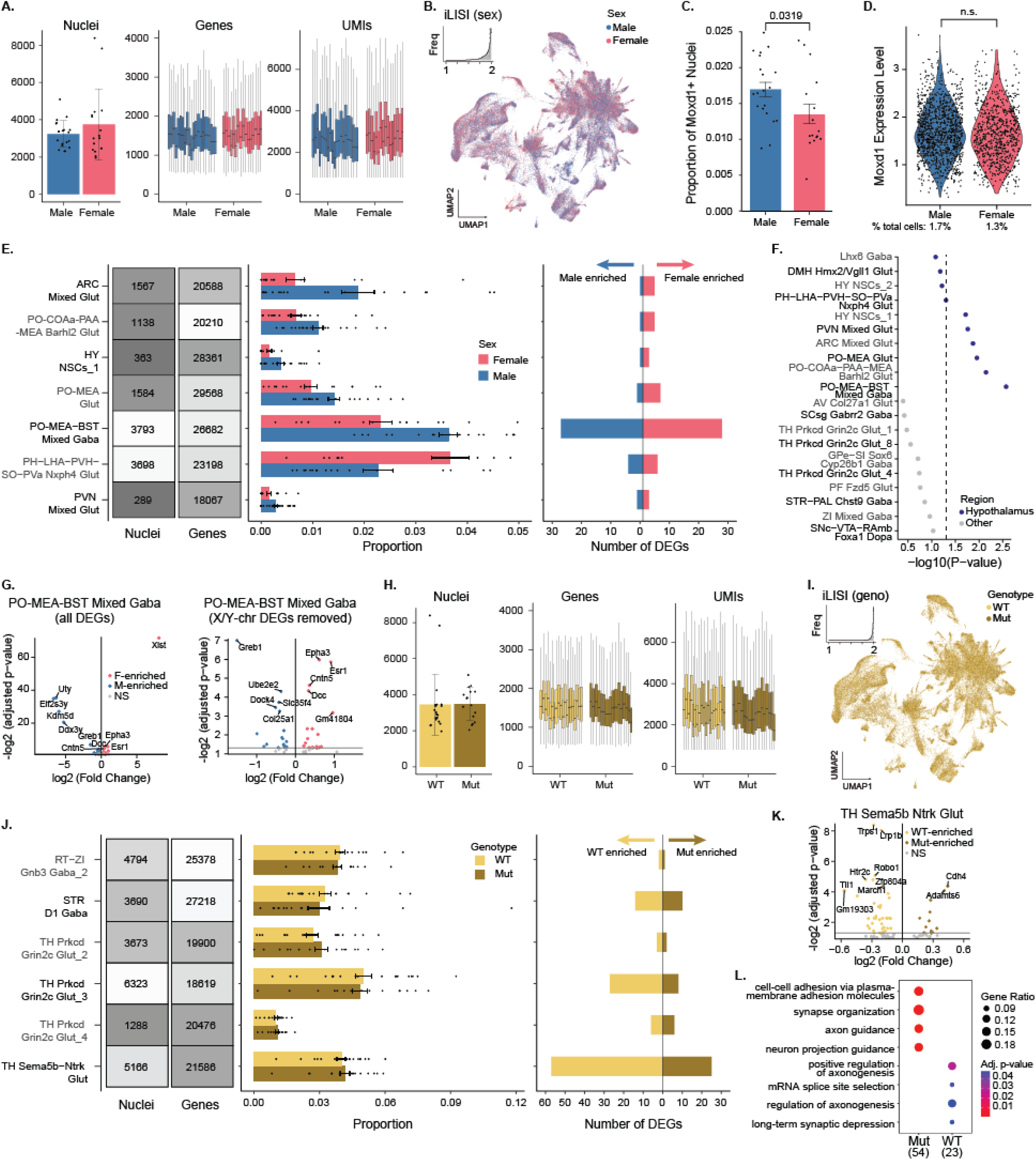
Hypothalamus shows sex and genotype differences in cell proportion and gene expression. **A.** General library statistics showing mean ± SD nuclei per sex (n = 20 males, 16 females), median ± SD genes per nucleus, and median ± SD unique molecular identifiers (UMIs) per nucleus. Box plots show the median (center line), interquartile range (box), and whiskers extending to 1.5 times the interquartile range. **B.** Uniform manifold approximation and projection (UMAP) showing 124,601 nuclei from the tuHY/TH colored by sex. Inset: Histogram showing the local inverse Simpson’s index (LISI) score with a median of 1.91, indicating that the sexes are well mixed and integrated. **C.** Bar plot showing mean ± SD proportion of *Moxd1*+ nuclei by sex, with a significant increase in males (p = 0.0319; linear regression). Points represent the mean expression of *Moxd1*+ neurons per sample. **D.** Violin plots of *Moxd1* gene expression across male and female samples, only including nuclei that have *Moxd1*+ expression > 0 (1.7% of male nuclei and 1.3% of female nuclei), with points representing each nucleus’ *Moxd1* levels. Differences not significant (p = 0.06744; Wilcoxon). **E.** From left to right: summary plot showing the numbers of nuclei and genes detected in each cluster; bar plot displaying the mean ± SEM relative proportions of nuclei in each annotated cell cluster for female and male samples; and number of differentially expressed genes (DEGs) per cell type that are upregulated in females (pink; n = 16 biological replicates) and upregulated in males (blue, n = 20 biological replicates) (*FDR adjusted p < 0.1, moderated t-test). **F.** Dot plot showing statistical significance (-log10(P-value)) of sex differences in cell type proportions, with hypothalamic regions (indigo) exhibiting more significant differences than other brain regions (gray) (p = 0.03271; Fisher’s) **G.** Volcano plot of DEGs by sex from PO-MEA-BST Mixed Gaba cluster, with female-enriched DEGs (F-enrich) in pink and male-enriched DEGs (M-enrich) in blue. DEGs that did not meet the p < 0.05 threshold but met the p < 0.1 threshold in gray (NS). Left plot includes all genes, right plot excludes X- and Y-chromosome genes for visualization. **H.** General library statistics showing mean ± SD nuclei per genotype (n = 20 *Myt1l* wild-type (WT), 16 *Myt1l* heterozygous mutant (Mut)), median ± SD genes per nucleus, and median ± SD UMIs per nucleus. Box plots show the median (center line), interquartile range (box), and whiskers extending to 1.5 times the interquartile range. **I.** UMAP showing the tuHY/TH colored by genotype. Inset: Histogram showing the LISI score with a median of 1.97, indicating that the genotypes are well mixed and integrated. **J.** From left to right: summary plot showing the numbers of nuclei and genes detected in each cluster; bar plot displaying the mean ± SEM relative proportions of nuclei in each annotated cell cluster for WT and Mut samples; and number of DEGs per cell type that are upregulated in WT (light yellow; n = 16 biological replicates) and upregulated in Mut (ochre, n = 20 biological replicates) (*FDR adjusted p < 0.1 moderated t-test). **K.** Volcano plot of DEGs from TH Sema5b Ntrk cluster, with wild-type-enriched DEGs (WT-enrich) in light yellow and mutant-enriched DEGs (Mut-enrich) in ochre. DEGs that did not meet the p < 0.05 threshold but met the p < 0.1 threshold in gray (NS). **L.** Gene Ontology (GO) analysis of differentially expressed genes between wild-type and mutant conditions, showing enriched biological processes related to neuronal development and function. Dot size indicates gene ratio and color intensity indicates significance.

A screen for further sex differences in cell-type proportions revealed several nominal differences (**Fig.2E)**. Male-biased clusters included arcuate glutamatergic neurons (ARC Mixed Glut) and multiple preoptic populations (PO-COAa-PAA-MEA Barhl2 Glut, PO-MEA Glut, PO-MEA-BST Mixed Gaba). In contrast, *Nxph4*+ glutamatergic neurons (PH-LHA-PVH-SO-PVa Nxph4 Glut) showed female-biased abundance. While none of these differences remained significant after multiple-comparison correction, the hypothalamus showed regional enrichment for sex differences overall: 6/37 hypothalamic clusters showed nominal sex differences compared to 0/28 non-hypothalamic clusters (Fisher’s exact test, *p*=0.033; **Fig2.F**). We then analyzed sex-biased gene expression within each subtype^33^, finding a total of 109 unique differentially expressed genes (DEGs) by sex. As expected, sex-chromosomal genes were differentially expressed across most clusters (**Fig2.E**). PO-MEA-associated neurons (PO-MEA-BST Mixed Gaba) exhibited the most DEGs, including the hormone receptor *Esr1* and membrane-associated proteins *Cntn5* and *Col25a1* (**Fig2.G**).

To ensure our findings generalize beyond the *Myt1l* context, we then analyzed WTs only. Key findings were recapitulated, including male-biased *Moxd1*+ neuron proportions and the prominence of PO-MEA-BST neurons in sex-differential gene expression (**ED-Fig2A-F**). Notably, sex differences in neuronal subtype proportions were more pronounced in WTs only, revealing significant sex dimorphism in neurons including preoptic, paraventricular, and ventromedial regions (**ED-Fig2D**)..

The effects of the loss of *Myt1l* were more subtle. We did not observe differences in neuronal proportions due to genotype after correction, with only two thalamic subtypes showing nominal differences (TH Prkcd Grin2c Glut and TH-PVT-RE Ntrk1-Nox4 Glut) (**Fig2.J**). However, DEG analysis revealed significant perturbations in neuronal subtype-specific gene expression (137 unique DEGs), with the largest number of DEGs observed in thalamic clusters (TH Prkcd Grin2c Glut_3 and TH Sema5b Ntrk Glut) (**Fig2.J**). DEGs included neurodevelopmental disorder-liability genes *Pten, Foxp2,* and *Nlgn1*^34^ that share features with MYT1L Syndrome, as well as ion channel genes *Kcnd2, Grik1, Cacna1c, and Gabrg3*, and were enriched for synaptic and axonal gene ontologies (**Fig.2K-L**), suggesting similarities to aberrant phenotypes in other regions of the brain observed following *Myt1l* loss^5,35^

### IEG activation identifies hypothalamic Agtr1a+ neurons as mediators of social reward seeking

Given the established use of IEGs as markers of neuronal activity at brain-wide and single-nucleus levels^10–18^, we analyzed IEG expression patterns across individual nuclei to identify neurons activated during the social operant paradigm. First, across the entire tuHY/TH, we identified significant increases in the canonical IEG *Fos*, marginal reductions in *Jun*, and no differences in *Arc* expression in non-achievers (**Fig.3A-C**), suggesting the presence of negative regulatory populations in the tuHY/TH. To thoroughly capture neuronal type-specific activation, we expanded beyond single IEGs and used a comprehensive list of IEGs^14^ to define “activated” nuclei (**Fig.3D**).

**Figure 3:**
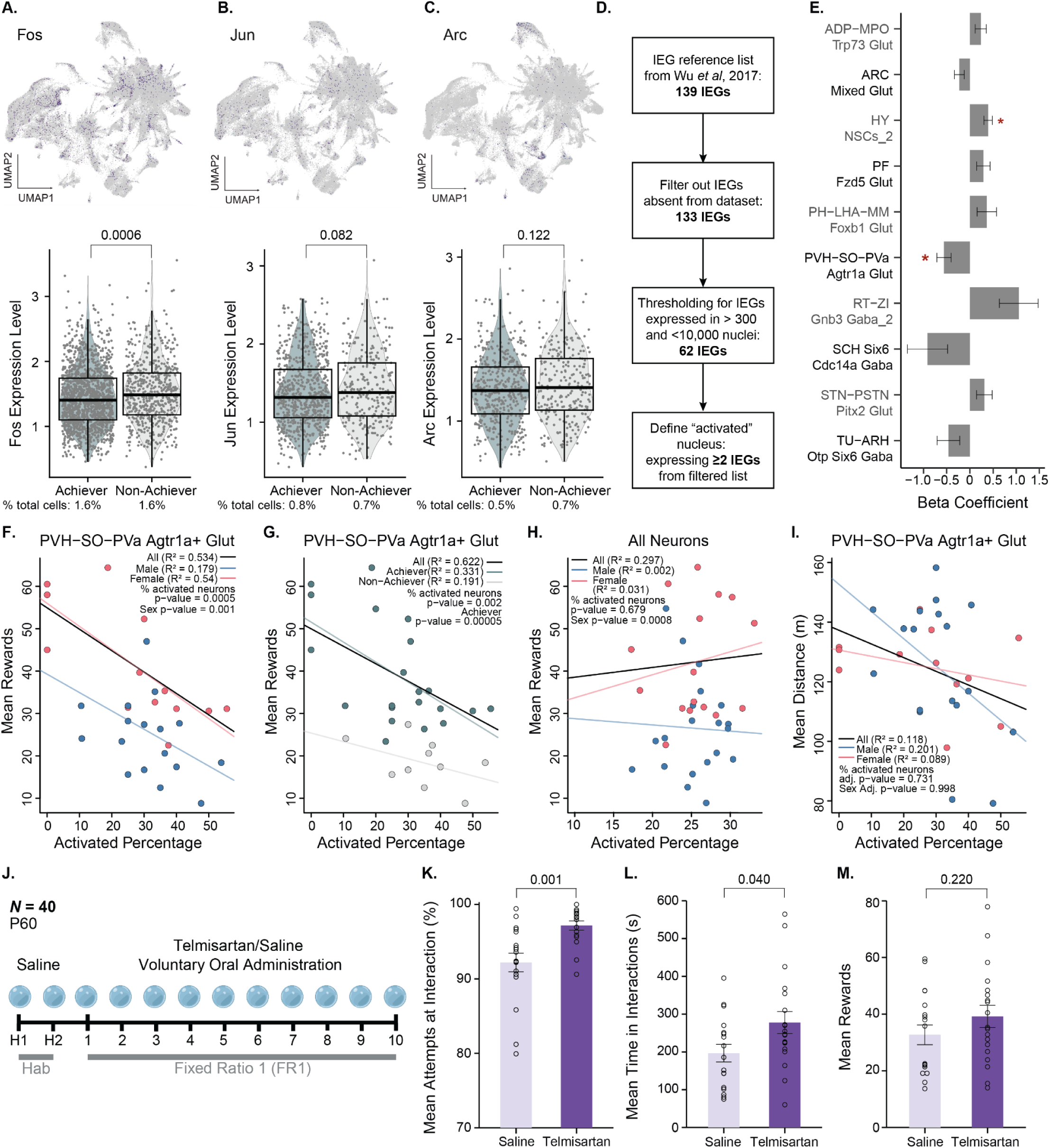
Neuronal activation identifies hypothalamic *Agtr1a* neurons as mediators of social orienting. **A-C**. UMAP plots showing expression of immediate early genes (IEGs) *Fos* (**A**), *Jun* (**B**), and *Arc* (**C**) across single nuclei, with corresponding violin plots, with box plots overlaid, below showing expression levels in neurons from Achiever versus Non-Achiever mice. Points represent the expression of each nucleus, and box plots represent the median (central line), inter-quartile range (box), and whiskers extending to 1.5 times the interquartile range. **D**. Flow chart describing the IEG filtering process for identifying activated neurons. Starting with a reference list of 139 IEGs from ^14^, 6 IEGs absent from the dataset were filtered out, leaving 133 IEGs. Further thresholding retained only IEGs expressed in >300 nuclei and <10,000 total nuclei, resulting in 62 final IEGs for analysis. “Activated” nuclei were then defined as those expressing ≥2 IEGs from this filtered list, creating a functional definition of neuronal activation for subsequent analyses. **E**. Bar plot showing beta coefficients from a linear regression analysis using percent activated nuclei for each neuronal cluster as a predictor of rewards in the social operant task. Only clusters with unadjusted p < 0.1 are shown, with error bars representing standard error. Asterisks indicate clusters that reached statistical significance after adjustment (adjusted p < 0.05), particularly highlighting the strong predictive relationship between PVH-SO-PVa *Agtr1a+* glutamatergic neuron activation and reward outcomes. **F-I**. Linear regression analyses examining relationships between neuronal activation and behavioral outcomes: **F**. Plot showing a negative relationship between mean rewards and percentage of activated PVH-SO-PVa *Agtr1a*+ glutamatergic neurons, with distinct patterns by sex (males: R² = 0.172; females: R² = 0.54). P-values of linear regression, including percent activated neurons (p = 0.0005) and sex (p = 0.001), indicate both are significant predictors. **G**. Similar analysis comparing Achiever versus Non-Achiever animals (Achiever: R² = 0.331; Non-Achiever: R² = 0.191). P-values of linear regression, including activated neuron percentage (p = 0.002) and achiever status (p = 0.00005), demonstrate a strong significance of both predictors. **H**. Relationship between mean rewards and activated percentage for all neurons across sexes, showing weaker correlations (males: R² = 0.002; females: R² = 0.031). P-values of linear regression for activated neuron percentage (p = 0.679) and sex (p = 0.0008) indicate that only sex significantly predicts rewards when considering all neurons. **I**. Analysis of mean distance traveled versus activated percentage of PVH-SO-PVa *Agtr1a*+ glutamatergic neurons by sex (males: R² = 0.201; females: R² = 0.089). P-values of linear regression for activated neuron percentage (p = 0.731) and sex (p = 0.998) suggest neither factor significantly predicts travel distance. **J**. Experimental design schematic for social operant testing showing telmisartan/saline voluntary oral administration protocol (*N* = 40), with habituation period (Hab; days H1 and H2) followed by fixed ratio testing (days 1-10). **K-M**. Comparison of behavioral outcomes between saline and telmisartan groups showing telmisartan administration significantly increases social orienting, as measured by attempts at interaction (defined as experimental animal entries into the investigation zone during a reward, p = 0.001; ANOVA) (**K**) and mean time spent in interactions (p = 0.040; ANOVA) (**L**), with no effect on mean rewards (p = 0.220; ANOVA) (**M**).

First, we compared the percent activation of each cluster between achievers and non-achievers. We found significant differences in two thalamic clusters, which largely overlapped with differences in activation by sex (**ED-Fig.3.A-B**). Therefore, to overcome potential sex-related sampling bias (given most females were achievers) and leverage the full individual variation in social motivation, we performed linear regressions for the percent activated neurons in each cluster and an animal’s mean rewards obtained (**Fig3.E**). Activity in *Agtr1a*+ neurons was negatively associated with rewards obtained on average in the task (**Fig.3F**). We confirmed this was true in the subset of data from *Myt1l* wild-type mice only, and repeated the regression, first including sex as a factor and then including achiever status as a factor, confirming that the negative correlation between *Agtr1a+* neuron activity and social rewards remained (**Fig.3G**, **Supplemental-Tables 5-6**). This finding was consistent whether we used the mean number of rewards obtained across all days or the number of rewards on the day of sacrifice (**Supplemental-Table 5**). As a control, we performed linear regressions using the mean activation of all neurons, not just *Agtr1a+* neurons, as a predictor of rewards and found no association (**Fig.3H**). Orthogonal analysis using Seurat’s gene module scores for the curated list of IEGs also highlighted paraventricular *Agtr1a*+ neurons as a mediator of social reward-seeking (**ED-Fig.3C-E, Supplemental-Table 5**). One possibility is that activity in *Agtr1a+* neurons simply reflects locomotion, as seeking rewards involves movement. Therefore, we repeated the regression to predict distance traveled rather than rewards. We found no clusters that significantly associated with locomotion in the task (**Fig3I, Supplemental-Table 5**).

Given the association between *Agtr1a*+ neuron activity and reduced social reward-seeking, we hypothesized that pharmacological blockade of AGTR1A would enhance social motivation. *Agtr1a* codes for angiotensin receptor 1a, the target of widely available and safe antihypertensive angiotensin receptor blockers (ARBs)^29^. To test this hypothesis, we initially delivered an ARB, losartan, or saline via daily intraperitoneal injection and assessed mice in the social operant assay (**ED-Fig.4**). While losartan-treated non-achievers showed increased social orienting compared to saline controls (p = 0.072, p = 0.049, ANOVA; **ED-Fig3K-L**), the injection procedure itself appeared to impair task performance, with reduced nose-poking in saline controls relative to wild-type controls from Cohort 1 (p = 0.021; **ED-Fig.3.B**). Indeed, in a separate cohort, we confirmed that intraperitoneal saline injection alone significantly reduced social reward-seeking (**ED-Fig.5**), prompting us to seek an alternative delivery method. We, therefore, administered telmisartan, an orally-available ARB^6^, via voluntary intake of gelatin cubes^36^ (**Fig.3J**). While telmisartan non-significantly increased total rewards (*p*=0.220), it robustly elevated social orienting: telmisartan-treated mice attempted to interact more (*p*=0.001) and spent more time in interactions (*p*=0.040) than saline controls (**Fig.3K-M, ED-Fig.6**). Together, these findings identify hypothalamic *Agtr1a+* neurons as negative regulators of social reward-seeking and demonstrate that clinically-available ARBs can enhance social motivation^37^, highlighting a promising therapeutic target for social deficits.

**Figure 4:**
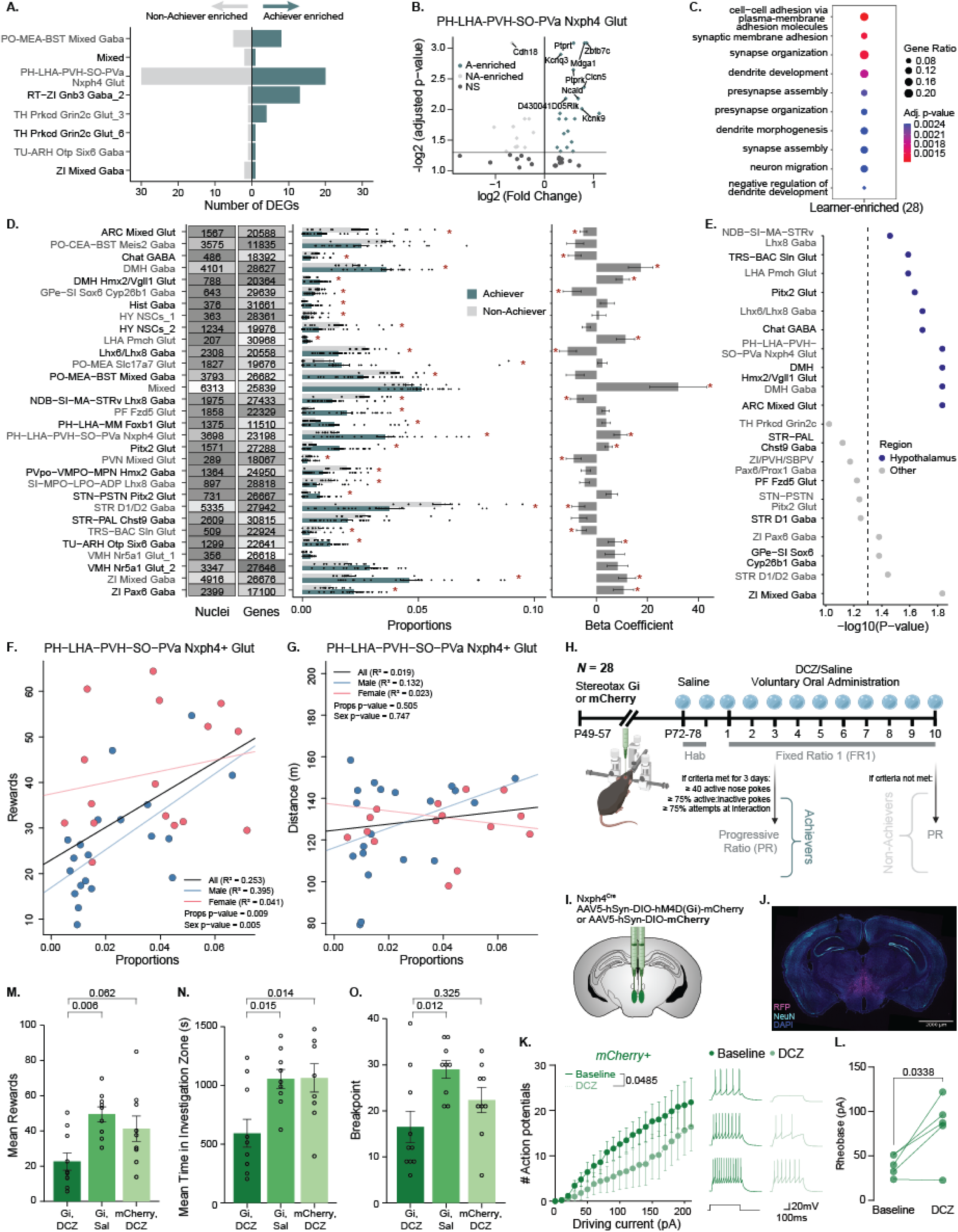
Variation in cell type proportions and degree of gene expression changes identify hypothalamic Nxph4+ neurons as mediators of social motivation. **A.** Bar plot showing the number of differentially expressed genes (DEGs) per cell type that are upregulated in Achievers (teal; positive values; n = 20) and upregulated in Non-Achievers (gray; negative values, n = 16) (*FDR adjusted p < 0.1, moderated t-test). **B.** Volcano plot of DEGs in PH-LHA-PVH-SO-PVa *Nxph4+* glutamatergic neurons, depicting log2(fold change) versus -log2(adjusted p-value). A-enriched genes (Achiever-enriched) are shown in teal, NA-enriched (Non-Achiever-enriched) in gray, and non-significant (NS) differentially expressed genes in dark gray. Top 10 most signifcant genes are labeled. **C.** Dot plot showing gene ontology enrichment analysis of achiever-enriched genes (28), with significant biological processes on the y-axis. Dot size represents gene ratio, and color indicates adjusted p-value significance. **D.** From left to right: summary plot showing the numbers of nuclei and genes detected in each neuronal cluster (clusters shown had significantly different proportions by achiever status before FDR correction OR significant findings in regression analysis); bar plots displaying the proportions across Achiever (teal) and Non-Achiever (gray) animals for each neuronal cluster (One sample (Het_F_5) removed for plotting for STR D1/D2 Gaba cluster); and bar plot showing Beta coefficients with error bars from regression analysis measuring the relationship between cell-type-specific proportions and social rewards. * indicates significant differences at p < 0.05 after FDR corrections. **E.** Dot plot showing statistical significance (-log10(P-value)) of linear regression analyses for rewards as predicted by neuronal proportions, with hypothalamic regions (indigo) exhibiting marginally more significant differences than other brain regions (gray) (p = 0.08719, Fisher’s). **F-G**. Linear regression analyses examining relationships between PH-LHA-PVH-SO-PVa *Nxph4*+ glutamatergic neuron proportions and behavioral outcomes: **F.** Plot showing a positive relationship between mean rewards and *Nxph4*+ neuronal proportions, with a stronger effect in males (male: R² = 0.395; female: R² = 0.041). P-values indicate a significant effect of proportions (p = 0.009) and sex (p = 0.005). **G.** Plot showing no significant relationship between mean distance traveled and Nxph4+ neuronal proportions (male: R² = 0.013; female: R² = 0.023; p=0.505). **H.** Experimental timeline for chemogenetic inhibition studies (N = 28), showing stereotaxic injection of viral constructs (P49-57) and subsequent testing paradigm with DCZ/saline administration via voluntary oral intake of gelatin cubes during fixed ratio 1 (FR1). **I.** Schematic showing stereotaxic delivery of AAV5-hSyn-DIO-hM3D(Gi)-mCherry or AAV5-hSyn-DIO-mCherry control virus to the lateral and posterior hypothalamus. **J.** Representative confocal mosaic reconstruction of immunohistochemistry verification of viral expression in the hypothalamus, with mCherry+ cells in magenta and NeuN+ cells in teal, and DAPI staining all nuclei. **K.** (Left) Input-output curve showing the number of action potentials elicited after a 600ms driving current during baseline and after 1uM DCZ in mCherry+ cells in the PH (n = 5 cells from 3 animals). p values indicate a significant effect of DCZ on excitability (p = 0.0485; two-way RM ANOVA). (Right) Representative current clamp traces at 50pA (top), 100pA (middle), and 150pA (bottom) during baseline and DCZ in the same cell. **L.** Rheobase of mCherry+ cells during baseline and after DCZ application (p = 0.0338, paired t-test). **M-O.** Behavioral outcomes following chemogenetic manipulation indicate inhibition of *Nxph4+* neurons in the posterior and lateral hypothalamus significantly reduces mean social rewards (p = 0.006; ANOVA), mean time in investigation zone attempting social interactions (p = 0.006; ANOVA), and overall social motivation, as measured by breakpoint, or the maximum effort exerted for a social interaction (p = 0.016; ANOVA).

### Variation in cell type proportions and degree of gene expression changes identify hypothalamic Nxph4+ neurons as mediators of social motivation

We next hypothesized that either stochastic variation in the properties of key cell types or variation in the proportion of key cell types might predict individual level variation in social motivation^1,7,19,21–23^. For the first possibility, we examined differential gene expression within specific neuronal populations. Ninety-six genes were differentially regulated between achievers and non-achievers (**Fig.4A**), with *Nxph4*+ neurons accounting for over 50% DEGs in this cluster included autism-associated genes (*Auts2*, *Megf11*)^38,39^, as well as neuronal ion channels (*Kcnm1*a*, Kcnq3*; **Fig.4B**). Gene ontology revealed enrichment for neurotransmitter- and behavior-associated functions (**Fig.4C**). Notably, sex and *Myt1l* genotype effects were larger than achievement-associated changes suggesting that natural variation in social motivation is only subtly associated with gene expression.

Next, we tested whether individual variation in social motivation is predicted by natural differences in tuHY/TH neuronal composition. First, comparison of neuronal subtype proportions between achievers and non-achievers revealed 26 significantly different neuronal populations, with more than two-thirds being hypothalamic, and 23 remained significant after accounting for sex in the model (**Fig.4D**). Next, to better account for the sex bias in achiever status and leverage the full natural variation of the data, we used linear regression to identify neuronal subtype proportions that predicted reward-seeking behavior. Since our findings depended on precise measurement of cellular compositions, we first verified the absence of experimenter-related variation (**ED-Fig.7A**). We then identified 17 neuronal subtypes significantly associated with mean rewards after FDR correction, which were marginally more likely to be hypothalamic (**Fig.4D-E**; **ED-Fig.8**). Among these, *Nxph4*+ glutamatergic neurons—which also showed the highest number of DEGs by achiever status—exhibited a significant positive correlation with reward-seeking (**Fig.4F**), independent of sex effects (**Supplemental-Table 7**). This correlation was reproducible in data from *Myt1l* wild-type animals only (**Supplemental-Table 7**). As a control, we evaluated whether neuronal proportions were predictors of overall distance traveled and found that none were significantly associated after FDR correction (**Fig.4G**; **ED-Fig.7B**), indicating that the tuHY/TH neurons sampled are more robustly linked to social reward than to locomotion.

Hence, we hypothesized that inhibiting *Nxph4*+ neurons would reduce social behavior. Sub-clustering analysis revealed that posterior and lateral hypothalamus (PH-LHA) *Nxph4*+ neurons drove the association with reward-seeking (**ED-Fig.9.A-B, Supplemental-Table 7**). Therefore, we targeted the PH-LHA of *Nxph4-Cre^mut/wt^*mice with inhibitory DREADD virus (hSyn-hM3D-Gi-mCherry) or control virus (hSyn-DIO-mCherry; **Fig.4H-I**)^8^. We confirmed successful targeting via immunofluorescence (**Fig.4J**) and functional inhibition upon agonist, deschloroclozapine (DCZ)^36^, application *ex vivo* (**Fig.4K-L, ED-Fig.10A-D**). After three weeks, we assessed social behavior following voluntary oral administration of DCZ or saline^36^. Inhibition of *Nxph4+* neurons in PH-LHA significantly reduced social reward-seeking, social orienting, and overall social motivation—maximum number of nose-pokes for a single reward—compared to both mCherry and saline controls (**Fig.4M-O, ED-Fig.10E-H**).

## DISCUSSION

Our findings provide proof-of-principle for a paradigm linking cellular mechanisms to behavioral variability across individuals. We demonstrate that individual variation in social motivation directly correlates with both the proportions and activity patterns of specific neuronal populations. The discovery that *Nxph4+* neuronal proportions positively predict social reward-seeking—confirmed through chemogenetic inhibition—provides compelling evidence that cellular composition contributes to behavioral variation. This supports a stochastic mechanism underlying individual differences, where developmental stochasticity crystallizes into fixed cellular architectures that deterministically shapes adult behavior—even among isogenic individuals^7,21–23^. This novel finding has potential implications beyond social motivation to other phenomena where individual variability is high in genetically identical individuals, like stress susceptibility^40^ and addiction vulnerability^41^. Thus, applying our approach to other brain regions and behaviors could identify additional cellular substrates of stable behavioral traits beyond social motivation.

Critically, our results reveal that no single cell type entirely predicts social motivation alone (model R^2^ < 0.31), underscoring the necessity of massively parallel approaches like single-cell transcriptomics to unravel the complex relationships between cells and behavior^19^. Single-cell combinatorial indexing platforms^3,4^ prove uniquely valuable for mapping individual variation, as they enable the sample sizes required to detect subtle yet significant correlations between cellular properties and behavior.

Our bird’s eye view and deep evaluation of the cell types of the tuberal hypothalamus revealed limited sexual dimorphism in neuronal proportions, with more pronounced differences in gene expression and neuronal activity—suggesting that functional rather than structural differences may drive sex effects on social motivation. A limitation of our approach is a broad regional sampling: deep-phenotyping analyses focused on specific nuclei of tuHY, like the ventromedial hypothalamus^18^, have successfully identified sex dimorphisms in proportions, indicating that more targeted approaches or even larger nuclei numbers may be better powered to detect structural sex difference.

For *Myt1l* mutation, we primarily observed gene expression changes rather than proportional differences, contrasting with a previous study in the cortex which found both^35^. This differential susceptibility to *Myt1l* mutation highlights the need for comprehensive analyses of a variety of brain structures in neurodevelopmental disorders. Finally, the identification of *Agtr1a+* neurons as negative regulators of social motivation is particularly promising, given that angiotensin receptor blockers are widely available medications with established safety profiles^6,29,37^. These findings could be immediately translatable for treating psychiatric conditions with social deficits^24–26^. Thus, future studies should explore whether AGTR1A antagonism enhances social motivation in preclinical models. Collectively, this work demonstrates how leveraging natural variation can reveal cellular substrates of complex behaviors, with direct implications for understanding and defining treatment targets for social impairments across neuropsychiatric conditions.

## Acknowledgements

We thank members of the Dougherty lab for helpful discussions and feedback, as well as Dr. John Constantino and Benjamin Boros for critical reading and manuscript suggestions. Thanks to Anthony Fischer for assistance with transporting Cohort 1 mice, Dr. Lex Kravitz for providing 3D brain matrices for dissections, and Kyle Kniepkamp for 3D printing support. We appreciate the expertise and services of the DNA Sequencing and Innovation Lab (DSIL) and the Genome Technology Access Center at the McDonnell Genome Institute (GTAC@MGI). We thank Dominic Skinner and Fatjon Leti from Scale Bio for technical support with the methodological and computational pipeline. We are grateful to Justin Wang and the Kravitz lab for donating DREADDs viruses. Additionally, we thank Dr. Christelle Anaclet for sharing the detailed DCZ voluntary oral administration protocol and Dr. Zachary Knight and the Knight Lab for generously donating Nxph4-Cre^mut/wt^ breeders. This work was funded by grants from the National Institute of Mental Health (RF1MH126723 to J.D.D. and R.D.M.; R01MH124808 to JDD, SEM; R01DA058755 and R01DA056829 to M.C.C.) and Simons Foundation Autism Research Initiative (734069 to J.D.D). S.S. was supported in part by the Autism Science Foundation (22-007).

## Author Contributions

Project conceptualization: S.S., S.K.K., L.Z.F., S.E.M., M.C.C., R.D.M., J.D.D. Method development, experiments, and data collection: S.S., S.K.K, L.Z.FM J.T., D.J.K., D.S., S.M.C., G.M.R. Formal analysis: S.S., L.Z.F, J.T., J.J. Figures and data visualization: S.S., S.K.K., L.Z.F., J.T., J.J. Writing-original draft: S.S., S.K.K., L.Z.F., J.T., D.S., M.R.B, J.D.D. Writing-review and editing: S.S., S.K.K., L.Z.F., J.T., D.J.K, D.S., S.M.C., S.E.M, M.C.C, R.D.M., J.D.D. Project coordination: S.S., S.E.M., M.C.C., R.D.M., J.D.D. Funding acquisition: S.S., S.E.M., M.C.C, R.D.M., J.D.D.

## METHODS

### Animal husbandry

All animal studies were approved by the Institutional Animal Care and Use Committee. Housing was a 12-hour light cycle, controlled temperature and humidity, and *ad libitum* food and water. The C57BL/6-*Myt1l^em1Jdd^*/J^5^ (Jackson #036428) line was crossed to C57BL/6J mice to generate experimental cohorts, genotyped as described^5^. As *Myt1l* loss is lethal postnatally, tested genotypes were *Myt1l* wild-type (WT) and *Myt1l* heterozygous mutants (Hets). Nxph4-2a-Cre knock-in mice (Jackson #033354) were a generous gift from Dr. Zachary Knight, and were genotyped as described^8^. After weaning at P21, all animals were group-housed by sex and genotype and/or treatment, with 3-5 mice per cage. Overall, five cohorts were used:

1. Assessed by social operant task, followed by single-nucleus RNA sequencing, cohort 1 included 36 mice, split by sex and *Myt1l* genotype (10 male WT, 10 male Hets, 10 female WTs, 6 female Hets). For all cohorts, unless noted, novel, sex- and age-matched stimulus animals were C57BL/6J mice bred in-house for all cohorts.
2. This cohort evaluated the effects of intraperitoneally-injected angiotensin receptor blocker, losartan, on social motivation. It consisted of n=40 P46 C57BL/6J mice (Jackson Laboratories #000664), balanced for sex. Mice were acclimated to WashU housing for 2 weeks prior to the task.
3. The third cohort piloted the effects of intraperitoneal injection of saline on social operant, using n=10(5M, 5F) from our C57BL/6J colony.
4. The fourth cohort received voluntary oral administration of angiotensin receptor blocker, telmisartan, and assessment of social motivation. It consisted of 40 C57BL/6J (Jackson Laboratories #0000664), balanced for sex. Stimulus animals were also purchased from Jackson Laboratories (#0000664). Mice were acclimated to WashU housing facility for 4 weeks.
5. The fifth cohort evaluated chemogenetic Gi inhibition of *Nxph4+* neurons via voluntary oral administration of DCZ on social motivation, using N=28 experimental Nxph4-Cre^mut/wt^ mice, balanced for sex.

All genotypes were assessed before weaning, and again after tissue harvest.

### Social Operant Behavior Testing

Social motivation, specifically measures of social reward seeking and social orienting as defined below, was evaluated using a social operant task as described^2^. Briefly, experimental and stimulus mice were handled for 3 days prior to starting the task and the tails of mice were marked with a non-toxic, permanent marker regularly to easily distinguish mice during testing. Male and female cages were separated in the testing room to avoid olfactory cue influence on behavior. Testing orders were randomly counterbalanced for group across apparatuses and trials. Testing began around P60 (P57-95) for all animals. All tasks were run during the light phase in a chamber illuminated with 75-80 lux of red light. The operant paradigm comprised habituation and testing trials. Habituation consisted of a 30-minute trial on two consecutive days, during which the door remained opened, and the nose-poke holes were blocked by panels to prevent any nose-poking prior to training. Training days consisted of 1-hr trials during which the fixed ratio 1 (FR1) reinforcement schedule was used to reward the mouse with a 12 s social interaction opportunity following one correct nose-poke. During the reward period, additional correct nose-pokes did not result in another reward. Task achievement criteria were attaining: a) at least 40 total nose-pokes, b) ≥75% accuracy (3:1 active:inactive nose-poke ratio), and c) ≥75% of rewards including an attempt at social interaction (defined as experimental mouse entering the social interaction zone for at least 1 s of the reward). After three consecutive days of showing achievement of learning criteria resulted in the mouse attaining “achiever” status and moving on to sacrifice for cohorts 1 and 4, or moving on to progressive ratio (PR) testing for cohorts 2, 3, and 5. Ten days of FR1 without reaching three consecutive days of criteria resulted in “non-achiever” status and moving on to sacrifice or PR testing as above. PR testing included a one-day trial to determine breakpoint, or the maximum effort exerted for a social interaction. During PR testing, rewards became progressively more effortful to obtain, with three additional nose-pokes required for each subsequent reward (i.e., 1, 3, 6, etc.). Breakpoint was defined as the number of nose-pokes a mouse exerted for its last reward.

### Statistical Analysis for Behavioral Outcomes

Statistical analyses and graph plotting were performed using IBM SPSS Statistics (v.27). Prior to analyses, data was screened for missing values and fit for assumptions of Analysis of Variance (ANOVA). This included the Shapiro-Wilk test on z-score-transformed data and qqplot investigations for normality, Levene’s test for homogeneity of variance, and boxplot and z-score (±3.00) investigation for identification of influential outliers. For cohort 4, from our initial cohort of 40 animals, 3 subjects were excluded from the final analysis due to methodological considerations, resulting in a final sample size of 37 animals for statistical analyses. Other than that, outliers were removed on a by-variable basis when skewness testing and z-scores indicated influential outliers. Our first ANOVA models included fixed effects terms: Genotype, Sex, and GenotypeSex for cohort 1; Treatment, Sex, and TreatmentSex for cohorts 2, 3, and 4; and Group, Sex, and Group*Sex for cohort 5. Subsequent ANOVA iterations removed terms for non-significant interactions and main effects, unless the interaction of a main effect was significant, until the most parsimonious model was achieved. Simple main effects tests (e.g., *t*-tests) dissected significant interactions post hoc. Multiple pairwise comparisons were Bonferroni corrected. Figure schematics were generated using BioRender. All datasets are available from the corresponding author upon reasonable request. Summary of statistical analyses for behavioral is in **Supplemental Table 2**.

### Dissections

Within 30 minutes of task completion for cohort 1, the P54-P76 mice were anesthetized and brains were quickly extracted, meninges removed, and the hypothalamus was dissected using 3D printed matrices (<u>tinyurl.com/y52tajsz</u>) to allow for precise extraction of tuberal hypothalamus from A/P -0.5 mm to -2.6 mm, M/L -1.5mm and 1.5mm and DV of 4-6mm. Dissected hypothalamus were flash frozen in liquid nitrogen, and stored at -80°C.

### Nuclei isolation and fixation

Nuclei from mice from cohort 1 were isolated from flash frozen tissue. Samples were processed in 9 batches of 4, counterbalanced for sex and genotype. We followed our lab’s nuclei isolation protocol, which is available on protocols.io (dx.doi.org/10.17504/protocols.io.dm6gp9qjdvzp/v1). The nuclei were stored at −80 °C until all samples were fixed.

### Single-nucleus RNAseq library preparation and sequencing

Libraries were prepared from cohort 1 (10 male WT, 10 male Hets, 10 female WTs, 6 female Hets). Frozen fixed nuclei were thawed on ice and counted twice via hemocytometer. Using the ScaleBio Single Cell RNA Sequencing Kit v1.0 (Scale Biosciences 2020008), all 36 samples were prepared in one batch. Nuclei were loaded at 10,000 per well (2-3 wells/sample) to add RT barcodes as sample identifiers. Pooled nuclei were distributed across the 384-well plate for Ligation Barcode addition, then pooled again with 1600 nuclei per well for second strand synthesis. After tagmentation and indexing PCR, libraries were pooled (5μl each), cleaned with 0.8X SPRIselect beads (Beckman Coulter B23317), and fragment size/concentration quantified using High Sensitivity D5000 Screentape (Agilent) and NEBNext Library Quant Kit (New England Biolabs E7630S). Sequencing was performed on a Novaseq X Plus (Illumina; shared 25B flowcell) to a target depth of 20,000 reads per nucleus.

### Single-nucleus RNAseq data processing

Base calls were converted to fastq format and demultiplexed by Index1 barcode by GTAC@MGI for each sample. We combined and gzipped the files into one R1 and R2 file and used a perl script to create the combined I1 file. Fastq files were demultiplexed, combined, and processed using ScaleBio’s nextflow pipeline (nf v.23.04.4, ScaleBio Seq Suite: RNA Workflow v1.5-beta2; github.com/ScaleBio/ScaleRna/tree/v1.5.0-beta2). Filtered count matrices were imported using Seurat v4.3.0^42^ to create Seurat objects for downstream analyses.

Quality control removed multiplets (DoubletFinder v2.0.3)^43^ and filtered for nuclei with >800 UMIs, 300-7000 genes, and <1% mitochondrial reads. The final dataset contained 138,033 nuclei (median: 2530 UMIs, 1447 genes per nucleus). Count matrices were log-normalized (scaling factor: 10,000), and top 3,000 variable genes were identified using VST. PCA was performed on these features (100 PCs calculated, top 50 used). Dimensionality reduction used UMAP. Clustering was performed using the Louvain algorithm (resolution 0.01-2.4), with optimal resolution (2.0) determined by clustree v0.5.0^44^.

Cluster markers were identified using FindAllMarkers (logfc.threshold=0.5, min.pct=0.25) and visualized via heatmaps. Filtered raw count data was converted to h5ad format with Ensembl IDs using org.Mm.eg.db v.3.16.0 and analyzed using MapMyCells (RRID: SCR_024672) to align the nuclei with the 10X Whole Mouse Brain (CCN20230722) reference^30^. For orthogonal annotation, nuclei were also projected onto HypoMap using mapscvi v.0.02 (github.com/lsteuernagel/mapscvi)^16^. Mapping results–class and subclass annotations and bootstrapping probabilities from MapMyCells and projected labels and prediction probability from HypoMap–were integrated into the Seurat object metadata. Based on annotations, glial cells comprised only 10,577 nuclei (8.2%) of the dataset.

A neuronal-only subset was created for further analysis, (124,601 nuclei; median: 2660 UMIs, 1498 genes) and underwent reprocessing as described above. Re-clustering with 50 PCs at resolution 1.8 yielded 75 initial clusters. Marker genes were identified using FindAllMarkers as above. To refine our clustering, we performed hierarchical clustering and manually compared annotation results across MapMyCells, HypoMap, and FindAllMarkers marker genes. We merged 8 cluster pairs based on shared marker profiles and removed 2 glial clusters, resulting in 65 final neuronal clusters. Clusters were annotated following Allen Brain Institute taxonomy^30,45^: predicted location, marker genes (or “Mixed” when nonspecific), and neurotransmitter type (Glut/Gaba).

### Differential gene expression and Gene ontology enrichment analysis

Differential gene expression analysis was performed using DESeq2 v1.38.3^33^. Single-cell expression data were aggregated into pseudobulk samples by summing counts for each gene within each sample-cell type combination. For each analysis (achiever status, sex, and genotype comparisons), DESeqDataSetFromMatrix was used with appropriate design formulas (∼ Sac_Learner, ∼ sex, or ∼ genotype). A negative binomial model was fitted for each gene, and a Wald test was performed for pairwise comparisons. Log2 fold changes were shrunk using the adaptive shrinkage estimator (ashr). Genes with FDR-adjusted p-value < 0.1 were considered differentially expressed, while genes with FDR-adjusted p-value < 0.05 were considered significant for volcano plotting. Complete statistical results from all gene expression analyses are provided in **Supplemental Table 3**. Gene ontology (GO) enrichment analysis was conducted using clusterProfiler v4.6.2^46^ with the enrichGO function. Significantly differentially expressed genes (DEGs) were used as query genes and analyzed separately for up- and down-regulated genes. All expressed genes within a given cluster were used as background. P-values were adjusted using the Benjamini-Hochberg method. Redundant GO terms were filtered using compareCluster, and results were visualized with dot plots to identify enriched biological processes for each condition. All analysis was repeated using only wild-type samples.

### Statistical analysis of cell type proportions

Differential cell type proportion analysis was performed using propeller in speckle v0.99.7^47^. For group comparisons (genotype, sex, achiever status, and dissector), the propeller function was used directly, which calculates cell type proportions for each cluster across samples and performs statistical testing between experimental conditions using logit-transformed values. Predicted anatomical annotations, based on cell type assignment, were integrated to assess region-specific effects, with statistical significance of proportion differences evaluated using Fisher’s exact tests. For associations between cell type proportions and behavioral metrics (rewards and distance traveled), proportions were extracted using the getTransformedProps function, logit-transformed, and tested using linear regression models with and without controlling for sex and achiever status. Benjamini-Hochberg correction was applied to adjust for multiple comparisons in all analyses. All key associations were replicated in a subset of data containing only wild-type samples. Complete statistical results from all proportion analyses are provided in **Supplemental Tables 4** and **6**.

### Immediate early gene (IEG) analysis

To examine neuronal activation patterns, we analyzed IEG expression in single nuclei. Due to the activity-dependent nature of IEG expression and the inherent sparsity of snRNA-seq data, we employed a comprehensive approach. Starting with an extensive list of 145 IEGs^14^, we filtered genes based on detection frequency, excluding those expressed in fewer than 300 nuclei (too rare) or more than 10,000 nuclei (approximately 10% of the dataset, to exclude more widely or potentially ubiquitously expressed genes). This filtering resulted in a curated list of 62 IEGs. A nucleus was classified as “activated” if it expressed two or more IEGs from this curated list. We then calculated the percentage of activated cells for each neuronal cluster and compared activation states between experimental groups (genotypes, sexes, and achiever status) using Fisher’s exact tests with Benjamini-Hochberg correction for multiple comparisons. Next, we performed regression analyses to assess association between neuronal activation in specific clusters and behavioral metrics, including mean rewards, rewards on day of sacrifice, mean distance, and distance on day of sacrifice. As a control, we also performed regression analyses on the entire neuronal population to distinguish cluster-specific effects from global activation changes when examining the relationship between activation and rewards.

We also computed a continuous IEG module score for each nucleus using the AddModuleScore function in Seurat. Briefly, the average expression of the IEGs in each cell is calculated and then subtracted by the aggregated expression of randomly selected control gene sets. A linear regression model was used to calculate the correlation between IEG module scores and the average number of total rewards for each cell type. All key findings were replicated in a subset of data containing only wild-type samples. Complete statistical results from all IEG analyses are provided in **Supplemental Table 5**.

### Drug Treatments

For Cohort 2, mice received either 10 mg/kg losartan in saline or saline via intraperitoneal injection 30-45 minutes prior to behavioral testing. For cohort 3, animals received either saline injection or no injection as control, again 30-45 minutes prior to behavioral testing. For both cohorts, mice underwent a 5-day handling period consisting of 2 days of standard handling followed by 3 days of scruffing habituation, to minimize stress during the experimental phase.

For drug administration in Cohorts 4 and 5, we employed a voluntary oral administration method using palatable gelatin cubes as described previously^36^ to replace the more stressful intraperitoneal injections. The gelatin stock solution was prepared with molecular-grade water containing 20% saccharine (Splenda), 14% gelatin (original unflavored gelatin, Knox), food flavoring (strawberry flavor, Frontier Co-op, USA), and blue food coloring (Whole Foods, USA), which was mixed at 50°C until fully dissolved. For cohort 4, telmisartan (10 mg/kg) was dispersed in gelatin matrix at 2 mg/mL concentration, with adult male mice receiving 125 μL gelatin cubes containing 0.25 mg telmisartan and adult females receiving 100 μL gelatin cubes containing 0.2 mg telmisartan. Due to telmisartan’s temperature sensitivity, these gelatin cubes were prepared at 30°C and stored at room temperature. For cohort 5, DCZ (0.5 mg/kg) was dissolved in gelatin matrix using a 5 mg/mL stock solution, with adult males receiving 125 μL gelatin cubes containing 2.5 μL of DCZ stock and adult females receiving 100 μL gelatin cubes containing 2 μL of DCZ stock. DCZ gelatin cubes were prepared at 50°C and stored at 4°C for up to 96 hours. For both cohorts, mice underwent a 5-day habituation period to drug-free gelatin cubes prior to experimental testing, with food deprivation required only for the first exposure. During experiment days, mice were individually transferred to holding cages where they consumed the gelatin cubes containing either drug or saline vehicle within 1-5 minutes before being returned to their home cages, with drug administration occurring 30-45 minutes prior to the start of experimental testing.

### Stereotaxic injections for DREADDs chemogenetics

The Nxph4-Cre^mut/wt^ mice in cohort 5 were anesthetized in an induction chamber with 5% isoflurane (SomnoSuite® Low-Flow Anesthesia System) and held on 1.5-2% during surgery. After anesthetization, mice were secured in the stereotaxic apparatus (Kopf Instruments Model 942 Small Animal Stereotaxic Instrument) using ear bars. 5-10 mg/kg of Meloxicam in sterile sodium chloride and 0.5 mL of sterile sodium chloride solution were injected. The surgical area was disinfected with iodine and 70% ethanol. 360 nl of AAV5-hSyn-DIO-hM4D(Gi)-mCherry (diluted to ∼1 x 10^12^ viral particles per mL, Addgene: 44362-AAV5) or AAV5-hSyn-DIO-mCherry (diluted to ∼1 x 10^12^ viral particles per mL, Addgene: 50459-AAV5) was injected into PH/LHA (from bregma: ML +/-0.4 mm, AP -2.3 mm, DV - 4.9 mm). The virus was infused in 6 cycles of 60nL at a rate of 10 nl/sec. Injections were made using Drummond Scientific Nanoject III and Nanoject Glass Capillaries. The Nanoject was left in place for 2 minutes after infusion and then raised by 0.05 mm for 8 minutes. Vetbond Tissue Adhesives (3M) was used to close the wound and topical antibiotic ointment was applied. Post-operative mice were left to recover on a heating pad with access to food and water.

### Perfusions, Sectioning, and Immunofluorescence

Mice in cohort 5 were perfused with ice-cold 1x PBS followed by 4% paraformaldehyde (PFA). Brains were harvested and postfixed overnight in 4% PFA, transferred to 15% sucrose solution for 24 h, then to 30% sucrose solution for another 24 h, frozen in OCT compound (Thermo Fisher, 23-730-571).

For IF validation, brains were cryosectioned to 35 um (Leica CM1950). Sections were blocked in 5% Normal Donkey Serum (Jackson Immunoresearch) in 1X PBS + 0.1% Triton-X for 1 h at room temperature. Primary antibodies were incubated overnight at 4°C (Rb @ RFP, 1:1000 (Rockland, 600-401-379); ms @ NeuN biotin conjugated, 1:100 (Sigma-Aldrich, MAB377B)). Alexa Fluor secondary antibodies (Dk @ Rb 568, 1:1000 (Invitrogen, A10042); Streptavidin 647, 1:1000 (Invitrogen, S32357)) were incubated for 1 h at room temperature. Nuclei were counterstained with DAPI at a 1:20,000 dilution for 10 min, washed, then mounted with Prolong Gold anti-fade mounting medium (Thermo Fisher, P36934). Confocal microscopy was performed on a STELLARIS 5 Confocal Microscope (Leica) with a 20x magnification lens.

### Perforated Patch Clamp Electrophysiology

A subset of *Nxph4-Cre^mut/wt^* mice from Cohort 5 expressing hM4Di in the PH (n=4 mice) were used for *ex vivo* electrophysiological validation of the DREADD after behavioural testing. Mice were deeply anesthetized under isoflurane, sacrificed, and the brain removed rapidly. 210um thick slices through the PH were made using a vibratome (Leica VT1200S) in an ice-cold choline chloride-based cutting solution (composition in mM: 0.5 CaCl2, 110 C5H14CINO, 25 C6H12O6, 25 NaHCO3, 7 MgCl2, 11.6 C6H8O6, 3.1 C3H3NaO3, 2.5 KCl and 1.25 NaH2PO4), bubbled continuously with 95 % O2 and 5 % CO2. Slices were incubated at 32oC for 15 minutes in artificial cerebrospinal fluid (ACSF; composition in mM: 119 NaCl, 2.5 KCl, 1.3 MgCl2, 2.5 CaCl2, 1.0 Na2HPO4, 26.2 NaHCO3, and 11 glucose) prior to moving to room temperature where they recovered for at least 30 minutes prior to recording.

For perforated patch recordings, hemi-sected slices were moved to a recording chamber perfused with ACSF containing 2 mM kynurenic acid and 100 uM picrotoxin maintained at 30oC (+/- 2oC). Neurons were visualized under infrared-differential interference contrast on an Olympus x56 upright microscope. LED illumination (CoolLED; 560nm) was used to identify cells expressing mCherry (hM4Di+) or cells not expressing mCherry (hM4Di-) in the PH. Borosilicate glass pipettes (3–5MΩ resistance) were pulled on a micropipette puller (Narishige PC-100) and filled with a K-gluconate-based solution (composition in mM: 130 K-gluconate, 10 creatine phosphate, 4 MgCl2, 3.4 Na2ATP, 0.1 Na3GTP, 1.1 EGTA, and 5 HEPES). Gramicidin (25ug/mL) was added to the internal solution prior to recording to allow for recording in the perforated configuration. After establishing the perforated configuration (5-30 minutes, monitored by responses to a -10 mV current injection), intrinsic properties of fluorescent and non-fluorescent cells were recorded during baseline conditions and after a 5 minute perfusion of 1uM DCZ through the recording chamber. Signals were amplified, filtered at 2 kHz, and digitized at 10 kHz using a MultiClamp 700B amplifier and Digidata 1550 (Molecular Devices). pClamp 11.4 (Molecular Devices) was used for data acquisition and analysis of electrophysiological parameters.

### Statistical Analysis

No statistical methods were used to predetermine sample sizes. For snRNAseq experiments, investigators were not blinded to the genotypes; however, all samples were processed in parallel on the same plates. Detailed descriptions of statistical analyses for specific experiments are provided in their respective sections above.

### Data availability statement

All raw and processed sequencing data has been submitted to GEO. It is available at accession number GSE291295.

### Code availability statement

All code required to reproduce this analysis is publicly available on Github: github.com/Dougherty-Lab/socop-hyp-snRNAseq

---

**ED Figure 1:**
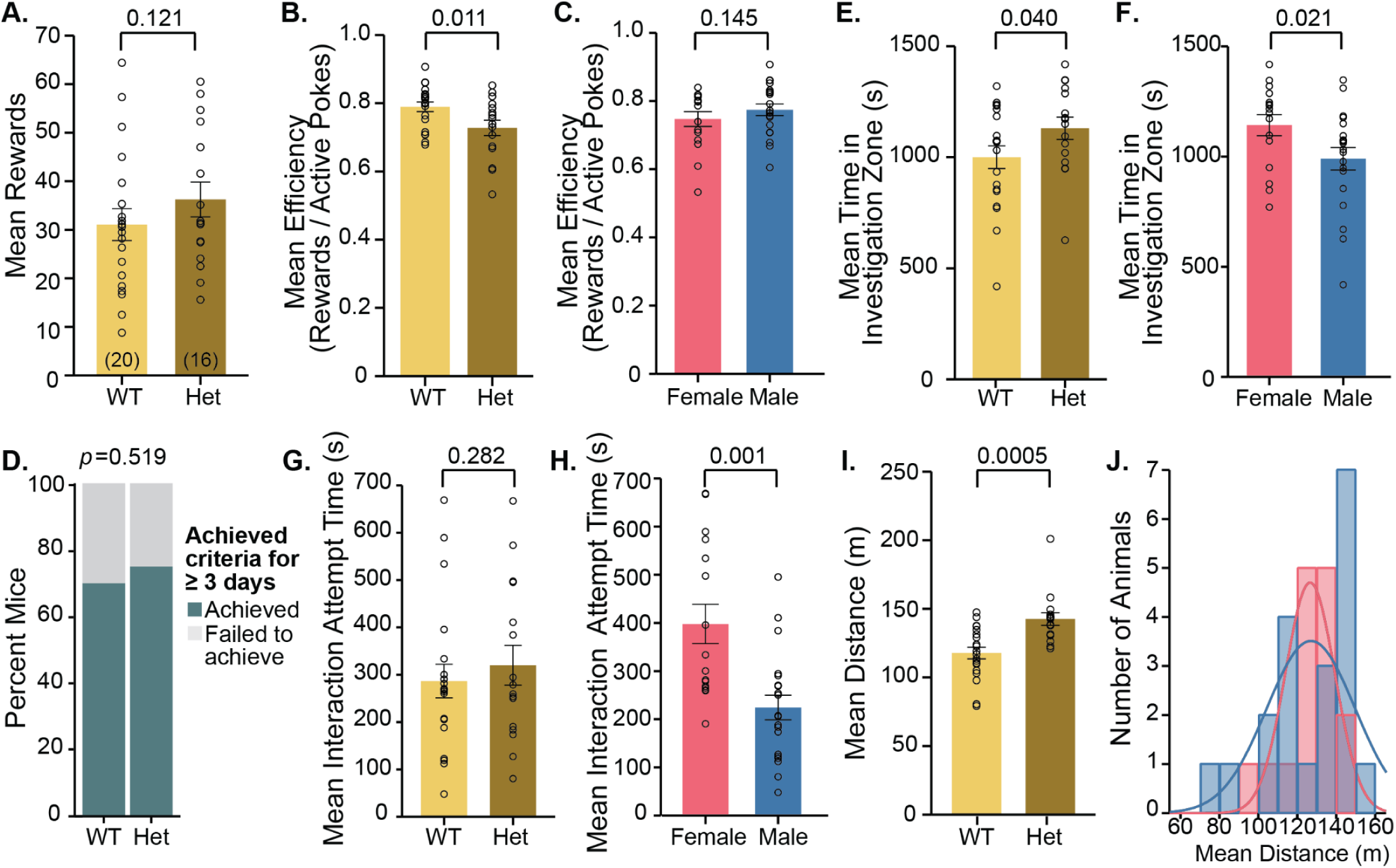
Mice display natural variation in social motivation that is influenced by sex but not genotype. **A-D.** Social reward-seeking variables: **A.** Mean rewards earned during social operant task comparing wild-type (WT, yellow) and heterozygous (Het, brown) mice (p=0.121; ANOVA). **B.** Mean efficiency (rewards/active pokes) comparing WT and Het mice (p=0.011; ANOVA). **C.** Mean efficiency split by sex (p=0.145; ANOVA). **D.** Percentage of mice achieving criteria for ≥3 days comparing WT and Het groups (left, p=0.519; Chi-squared). Criteria: ≥40 total nose-pokes, ≥75% accuracy, ≥75% attempts at interaction. **E-H.** Social orienting variables: **E.** Mean time spent in investigation zone (in seconds) comparing WT and Het mice (p=0.040; ANOVA). **F.** Mean time in the investigation by sex (p=0.021; ANOVA). **G.** Mean interaction attempt time (in seconds) comparing WT and Het mice (p=0.282; ANOVA). **H.** Mean interaction attempt time (s) split by females versus males (p=0.001; ANOVA). **I.** Mean distance traveled (m) comparing WT and Het mice (left, p=0.0005; ANOVA). **J**. Histogram showing distribution of mean distances across all animals, split by sex.

**ED Figure 2:**
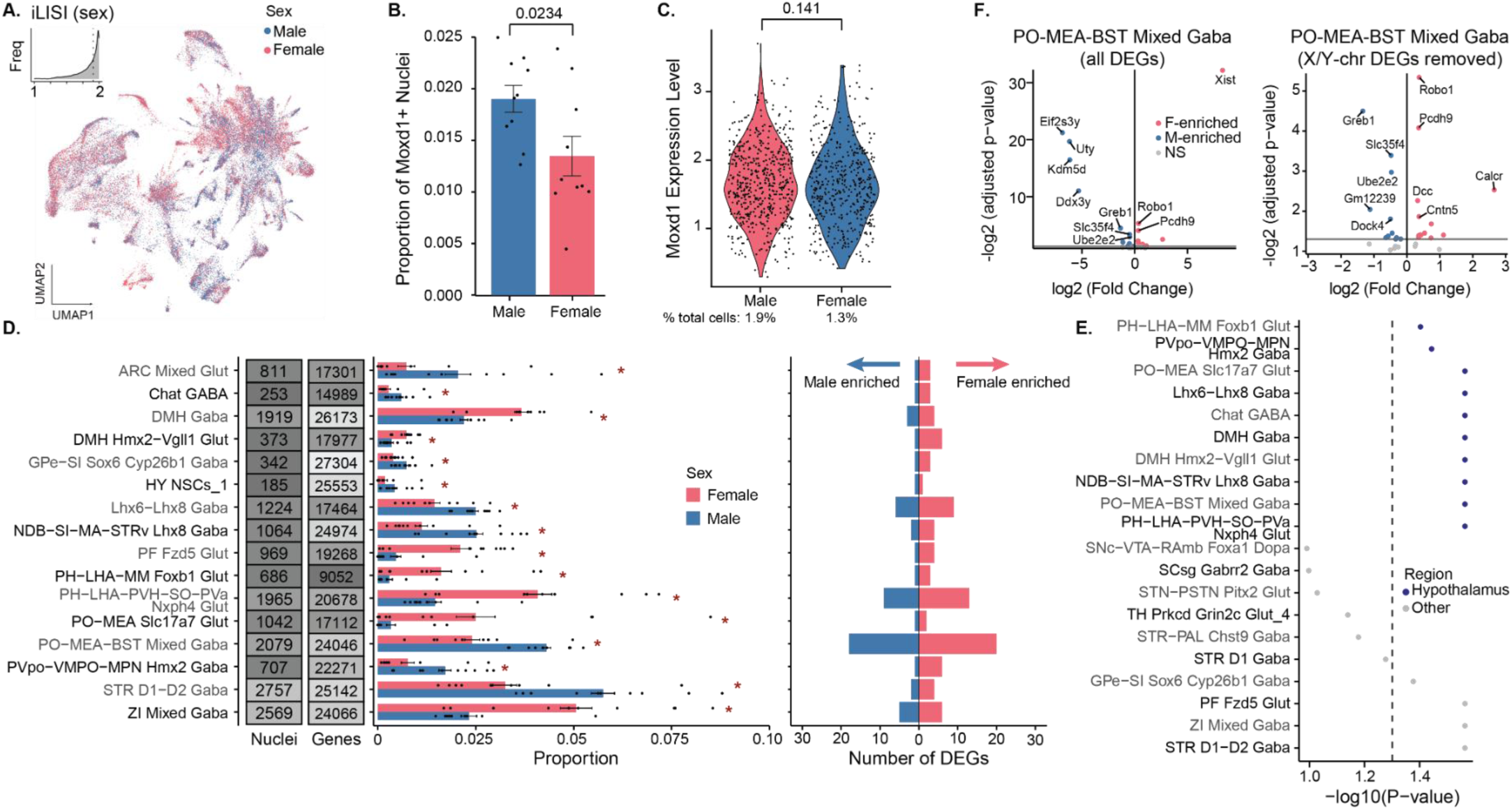
In wild-type (WT) mice only, hypothalamus shows more pronounced differences in cell proportion and some differences in gene expression. **A.** Uniform manifold approximation and projection (UMAP) showing 64,619 nuclei from the tuHY/TH from WT animals only, colored by sex. Inset: Histogram showing the local inverse Simpson’s index (LISI) score with a median of 1.84, indicating that the sexes are well mixed and integrated. **B.** Bar plot showing mean ± SD proportion of *Moxd1*+ nuclei by sex, with a significant increase in males (p = 0.0234; linear regression). Points represent the mean expression of *Moxd1*+ neurons per sample. **C.** Violin plots of *Moxd1* gene expression across male and female samples, only including nuclei that have *Moxd1*+ expression > 0 (1.9% of male nuclei and 1.3% of female nuclei), with points representing each nucleus’ *Moxd1* levels. Differences not significant (p = 0.141; Wilcoxon). **D.** From left to right: summary plot showing the numbers of nuclei and genes detected in each cluster; bar plot displaying the mean ± SEM relative proportions of nuclei in each annotated cell cluster for female and male WT samples; and number of differentially expressed genes (DEGs) per cell type that are upregulated in WT females (pink; n = 10 biological replicates) and upregulated in WT males (blue, n = 10 biological replicates) (*FDR adjusted p < 0.1, moderated t-test). **E.** Dot plot showing FDR-adjusted statistical significance (-log10(P-value)) of sex differences in cell type proportions, with hypothalamic regions (blue) exhibiting no more significant differences than other brain regions (gray) (p = 0.146; Fisher’s) **F.** Volcano plot of DEGs by sex from PO-MEA-BST Mixed Gaba cluster, with female-enriched DEGs (F-enrich) in pink and male-enriched DEGs (M-enrich) in blue for WT animals. DEGs that did not meet the p < 0.05 threshold but met the p < 0.1 threshold in gray (NS). Left plot includes all genes, right plot excludes X- and Y-chromosome genes for visualization purposes

**ED Figure 3:**
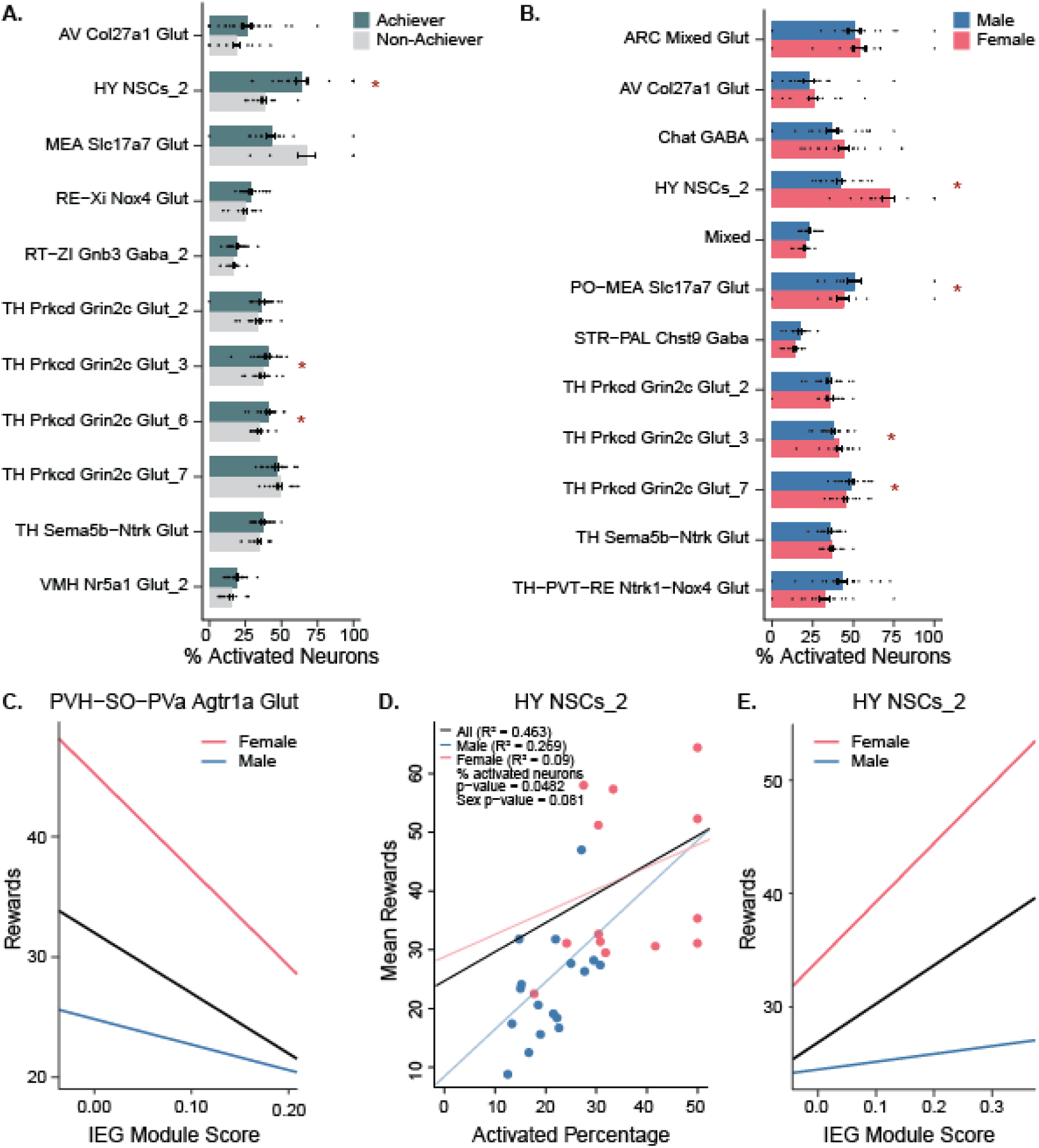
Module-based analysis of Immediate Early Gene activation also identifies Agtr1a neurons. **A.** Bar plot showing percent activated neurons as determined by filtered IEG expression across each neuronal cluster for Achievers (teal) and Non-Achievers (gray). **B.** Bar plot showing percent activated neurons across each neuronal cluster for Male (blue) and Female (pink) animals. Clusters shown were nominally significant before FDR corrections * indicates significant differences at p < 0.05 after FDR corrections. **C.** Using the Seurat AddModuleScore method to quantify immediate early gene (IEG) expression, which calculates a score that subtracts the average expression of a random control gene set from the average expression of curated IEG genes confirms the negative correlation observed in Figure 3F. Linear regression showing negative relationship between IEG expression score in PVH-SO-PVa Agtr1a+ glutamatergic neurons and mean rewards, with distinct patterns between females (green line, steeper negative slope) and males (black line). **D.** Scatterplot showing positive relationship between mean rewards and percentage of activated HY NSCs_2 neurons, with distinct patterns by sex (males: R² = 0.265; females: R² = 0.093). P-values of linear regression including percent activated neurons (p = 0.0482) and sex (p = 0.081) indicate both are significant predictors. **E.** Linear regression showing relationship between IEG expression score and mean rewards in HY NSCs_2 neurons as determined by module-based analysis, replicating the positive correlation in **B**.

**ED Figure 4:**
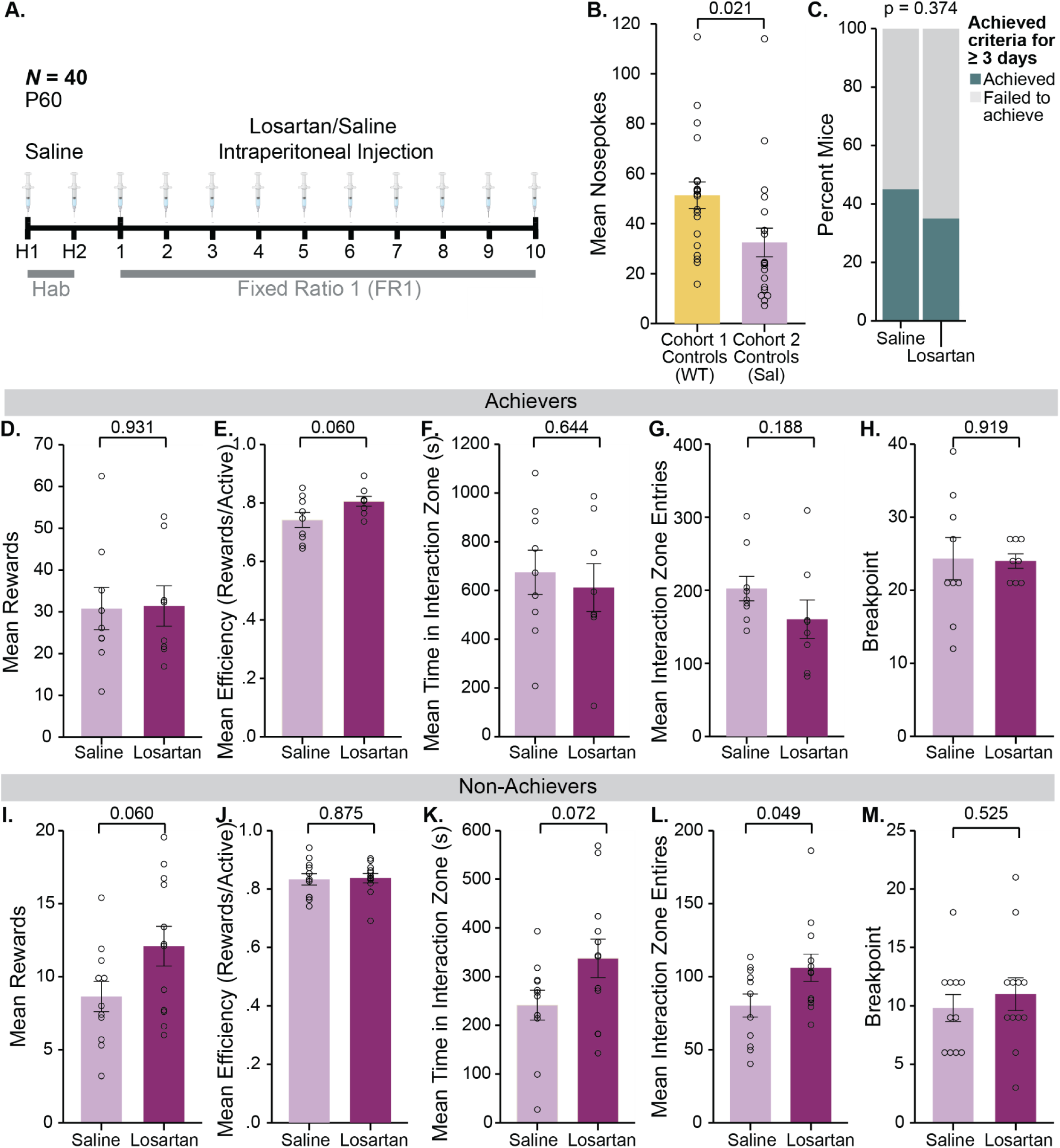
Losartan marginally increases social reward seeking and orienting in non-achievers. **A.** Schematic of angiotensin receptor blocker (ARB), losartan, intraperitoneal injection during social operant task. All mice received intraperitoneal injections of either saline (control) or losartan (ARB) 45 minutes before task initiation. **B.** Mean nose-pokes in wild-type animals from Cohort 1 and saline control animals from current cohort, Cohort 2, showing significantly lower nose-poking following intraperitoneal saline injection (p = 0.021). **C.** Percentage of achievers (teal) versus non-achievers (gray) in saline controls and ARB (losartan) treatment groups. **D-H** Results from achievers only: **D.** Mean rewards comparing saline and losartan treatment (p=0.931) **E.** Mean efficiency, defined as fraction of active nose-pokes rewarded (p=0.060) showing marginal effect of losartan, **F.** Mean investigation zone time by treatment (p=0.644), **G.** Mean investigation zone entries by treatment (p=0.188), **H.** Breakpoint, the maximum number of nose-pokes a mouse exerted for a reward, by treatment (p=0.826). **I-M.** Results from non-achievers only: **I.** Mean rewards (p=0.060) showing marginal increase with losartan treatment, **L.** Investigation zone time (p=0.072) with marginal increase in social orienting following losartan treatment, **J.** No effect on efficiency following losartan treatment (p = 0.875), **K.** Investigation zone entries (p=0.049) showing significant increase with losartan treatment, and **N.** Breakpoint (p=0.525) showing no significant difference between treatment groups.. Criteria for achiever status: ≥40 total nose-pokes, ≥75% accuracy, ≥75% attempts at interaction for ≥3 days. All comparisons were made by ANOVA.

**ED Figure 5:**
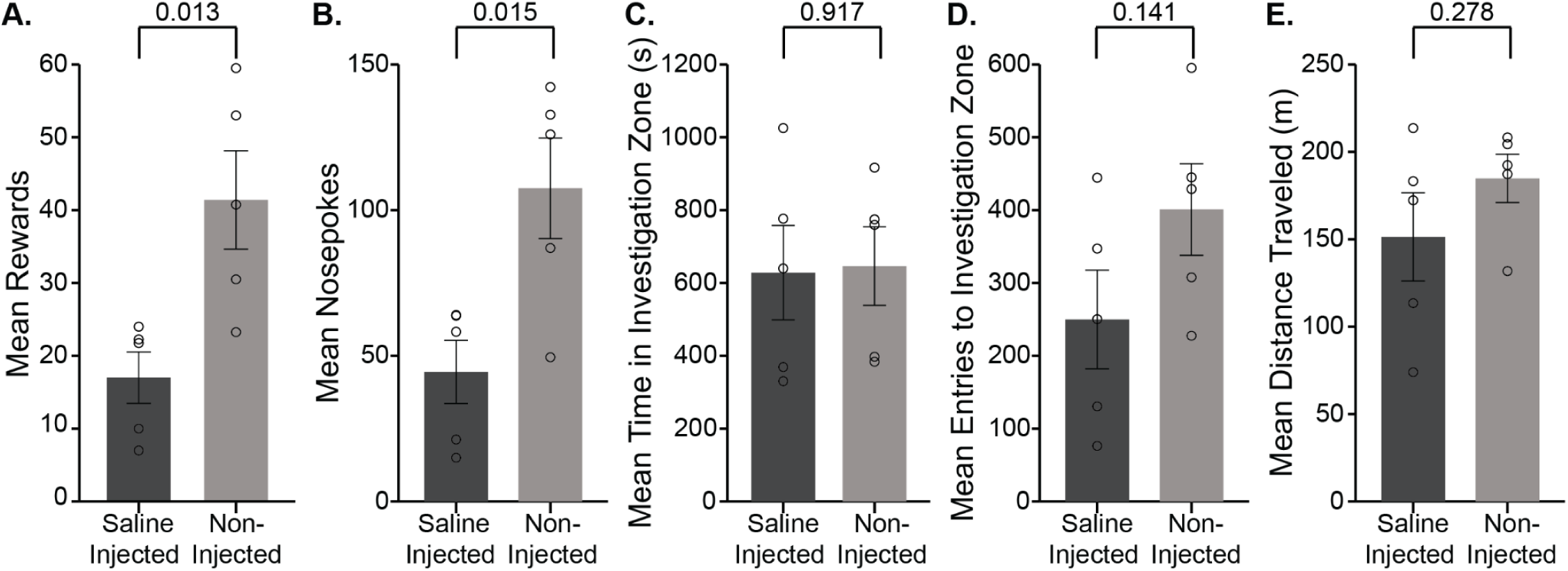
Intraperitoneal injection reduces performance in social operant task. **A.** Average total rewards earned during the social operant task comparing saline-injected (dark gray) and non-injected control mice (light gray) showing significant reduction in reward-seeking in injected mice (p=0.013). **B.** Average total nose-pokes showing significantly reduced task engagement in saline-injected mice compared to non-injected controls (p=0.015). **C.** Average exploratory zone time (in seconds) comparing saline-injected and non-injected mice, with no significant differences between groups (p=0.917). **D.** Average interaction zone entries comparing saline-injected and non-injected mice, showing no significant differences (p=0.141). **E.** Average test distance traveled (in m) comparing saline-injected and non-injected control mice, with no significant differences in locomotor activity (p=0.278). Data presented as mean ± SEM with individual data points shown. All comparisons were made by ANOVA.

**ED Figure 6:**
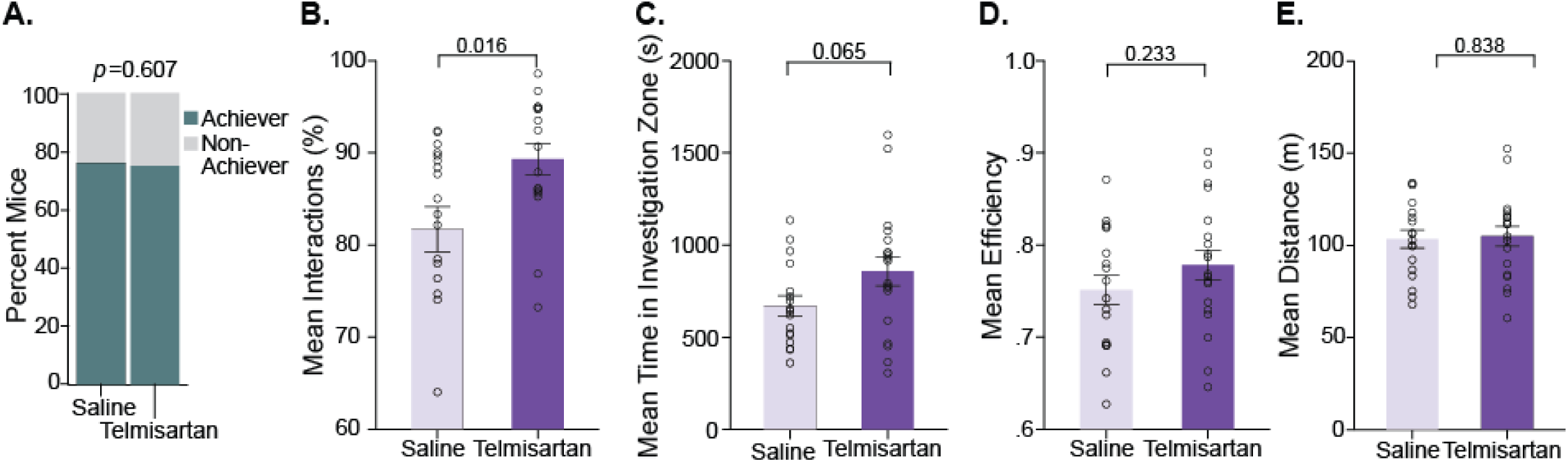
Telmisartan modulation specifically impacts social orienting but not reward-seeking. **A.** Percentage of mice achieving social reward-seeking criteria comparing Saline and Telmisartan groups, showing no significant difference (p=0.607), indicating telmisartan modulation does not affect ability to meet social reward-seeking criteria. **B.** Social orienting: Mean percent of rewards that contained an interaction (both experimental and stimulus animal in the investigation zone) comparing Saline and Telmisartan groups, showing significantly increased percent interactions in the Telmisartan group (p=0.016). **C.** Social orienting: Mean time spent in the investigation zone (in seconds) comparing Saline and Telmisartan groups, showing marginally significant increase in the Telmisartan group (p=0.065). **D.** Social reward-seeking: Mean efficiency (rewards/ active pokes) comparing Saline and Telmisartan groups, showing no significant difference (p=0.233), further confirming that telmisartan modulation does not impact reward-seeking behaviors. **E.** Control: Mean distance traveled (in mm) comparing Saline and Telmisartan groups, showing no significant difference (p=0.838), indicating that changes in social orienting measures are not due to general locomotor differences.

**ED Figure 7:**
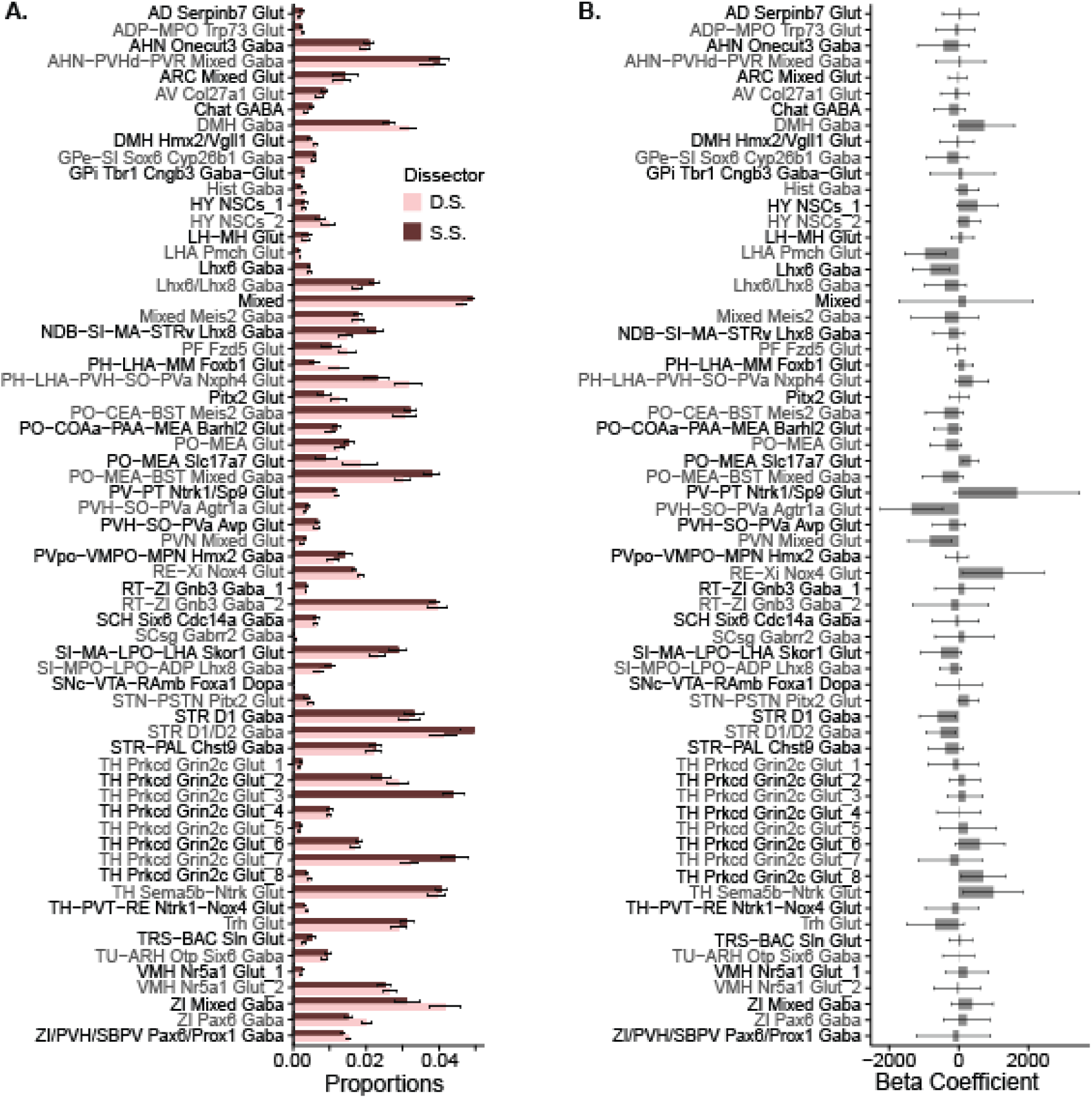
Controls for proportion analyses indicate no effect of dissector or locomotion. **A.** Bar chart of proportions by dissector (D.S. or S.S.), showing dissector does not significantly influence neuronal proportions in tuHY/TH, as seen by comparable proportions. **B.** Bar chart of beta coefficients for linear regressions of proportions as predictors of distance travelled, indicating no cell types had proportions that were a significant predictor of distance in the tuHY/TH.

**ED Figure 8:**
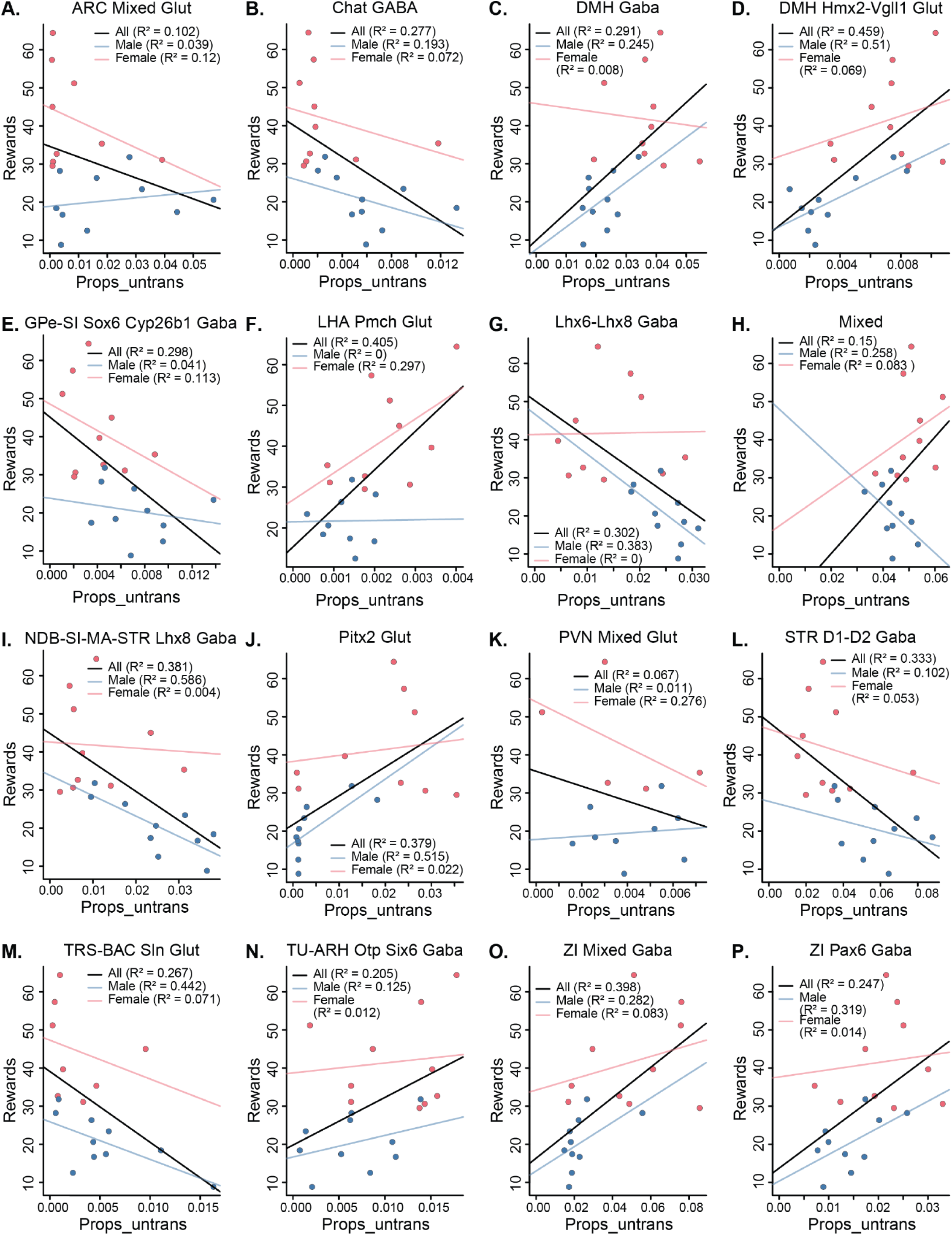
Additional cell types also associate with social motivation. **A-P.** Scatterplots for all clusters that had significant linear regression findings as predictors of social rewards in alphabetical order (with the exception of PH-LHA-PVH-SO-PVa Nxph4+ Glut which is shown in Fig.4F). Each plot includes R² values for all subjects, males, and females. Pink dots represent females and blue dots represent males, with corresponding regression lines showing the relationship between neuronal proportions (x-axis) and rewards (y-axis) for each sex. P-values can be found in **Supplemental-Table 7**.

**ED Figure 9:**
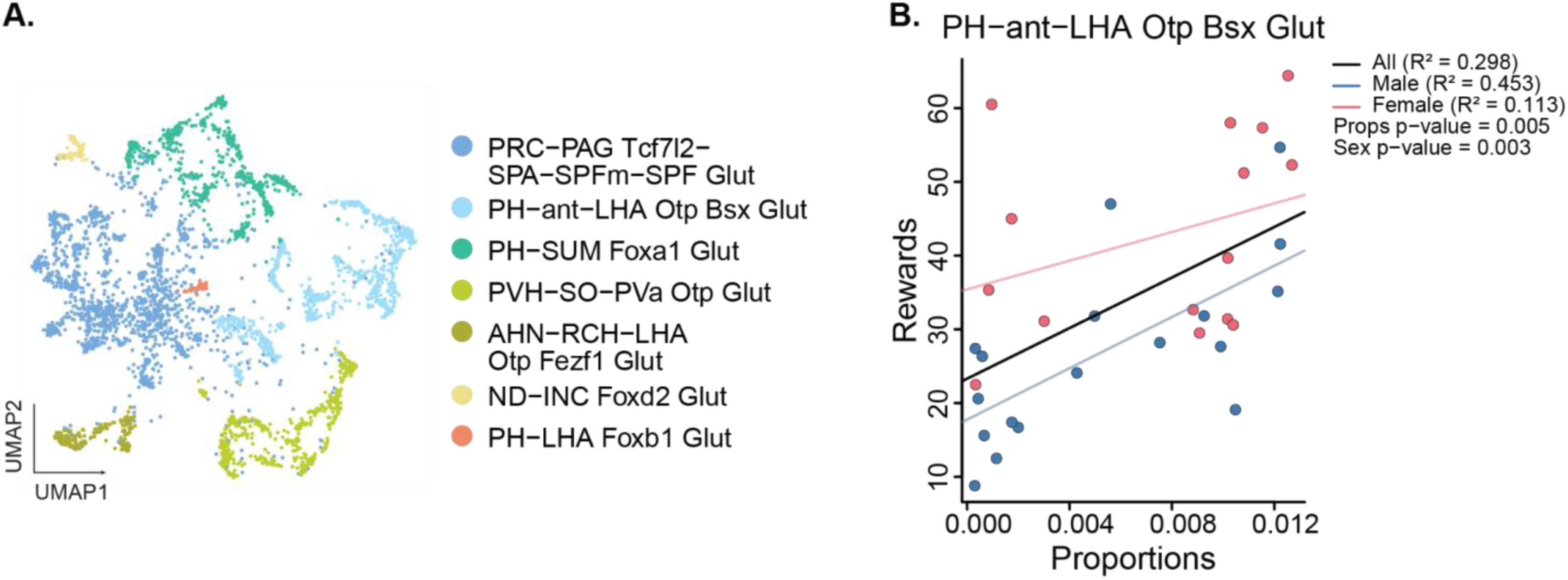
PH-LHA subset of *Nxph4+* neurons is associated with reward seeking. **A.** UMAP showing 8 distinct clusters obtained by subsetting PH-LHA-PVH-SO-PVa Nxph4+ glutamatergic neuronal cluster, roughly split by predicted region. **B.** Scatterplot for proportions of subcluster PH-ant-LHA Otp Bsx Glut showing a positive correlation with rewards even after correcting for sex P = 0.005), with similar directions in male and female (males: R² = 0.453; females: R² = 0.113).

**ED Figure 10:**
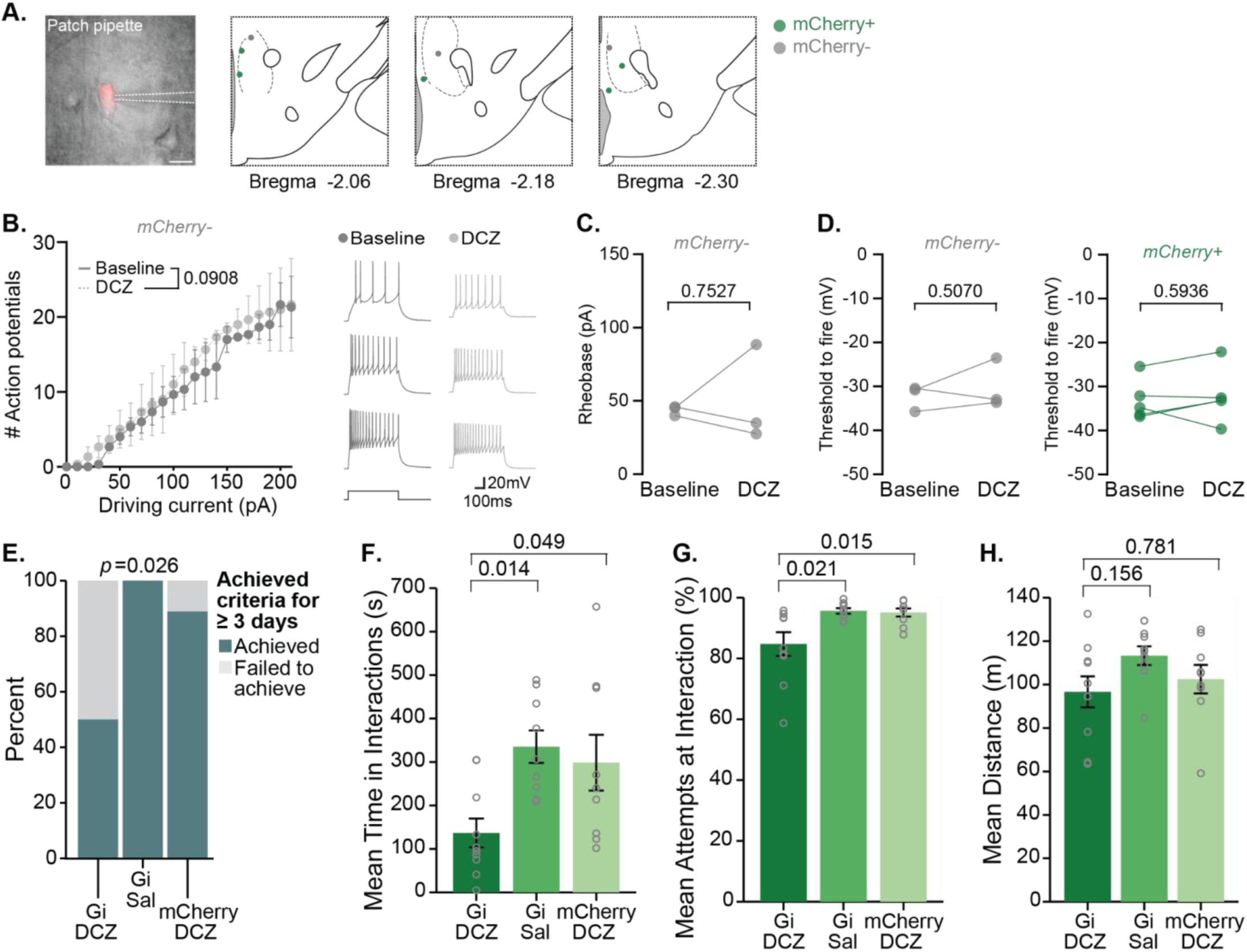
Nxph4 inhibition impacts multiple measures of social motivation. A. Representative images showing expression of mCherry+ virus at different stereotaxic coordinates (from - 2.06 to -2.30) in the posterior hypothalamus, with green labeling indicating mCherry expression. B. (Left) Input-output curve showing the number of action potentials elicited in response to increasing driving current (0-200 pA) during baseline and after DCZ administration in mCherry-cells. DCZ has no effect on the excitability of mCherry-cells (p = 0.0908, two-way RM ANOVA). (Right) Representative current clamp traces at 50pA (top), 100pA (middle), and 150pA (bottom) during baseline and DCZ in the same mCherry-cell.. C. Individual data points showing changes in rheobase (pA) following DCZ administration compared to baseline, with connecting lines representing paired measurements from the same cells in electrophysiological recordings. DCZ administration has no effect on rheobase of mCherry-cells (p = 0.7527, paired t-test). D. Individual data points showing changes in action potential threshold (in mV) following DCZ administration in both mCherry-control cells (left) and mCherry+ experimental cells (right), with connecting lines representing paired electrophysiological measurements from the same cell. DCZ administration does not affect the threshold to fire in either mCherry- (p = 0.5070, paired t-test) or mCherry+ cells (p = 0.5963, paired t-test). E. Bar graph showing the percentage of mice achieving social interaction criteria across three experimental groups (Gi_DCZ, Gi_Sal, and mCherry_DCZ), with significant differences observed (p=0.026, ANOVA). F. Mean time in interactions (in seconds) across the three experimental groups (Gi_DCZ, Gi_Sal, and mCherry_DCZ), showing significant reduction in the Gi_DCZ group (p=0.011, ANOVA). G. Mean percent successful daily interactions across the three experimental groups, showing significant differences with reduced interactions in the Gi_DCZ group (p=0.008, ANOVA). H. Mean distance traveled (in mm) across the three experimental groups, showing no significant differences (p=0.176, ANOVA), indicating that changes in social motivation measures are not due to general locomotor deficits

